# Structure and assembly of the diiron cofactor in the heme-oxygenase-like domain of the *N*-nitrosourea-producing enzyme SznF

**DOI:** 10.1101/2020.07.29.227702

**Authors:** Molly J. McBride, Sarah R. Pope, Kai Hu, Jeffrey W. Slater, C. Denise Okafor, Emily P. Balskus, J. Martin Bollinger, Amie K. Boal

## Abstract

In biosynthesis of the pancreatic cancer drug streptozotocin, the tri-domain nonheme-iron oxygenase, SznF, hydroxylates *N*^δ^ and *N*^ω^’ of *N*^ω^-methyl-L-arginine before oxidatively rearranging the triply modified guanidine to the *N*-methyl-*N*-nitrosourea pharmacophore. A previously published structure visualized the mono-iron cofactor in the enzyme’s C-terminal cupin domain, which effects the final rearrangement, but exhibited disorder and minimal metal occupancy in the site of the proposed diiron cofactor in the *N-*hydroxylating heme-oxygenase-like (HO-like) central domain. Here we leverage our recent report of an intensely absorbing *µ*-peroxodiiron(III/III) intermediate formed from the Fe_2_(II/II) complex and O_2_ to understand assembly of the diiron cofactor in the HO-like domain and to obtain structures with both SznF iron cofactors bound. Tight binding at one diiron subsite is associated with a conformational change, which is followed by weak binding at the second subsite and rapid capture of O_2_ by the Fe_2_(II/II) complex. Differences between iron-deficient and iron-replete structures reveal both the conformational change required to form the O_2_-reactive Fe_2_(II/II) complex and the structural basis for cofactor instability, showing that a ligand-harboring core helix dynamically refolds during metal acquisition and release. The cofactor also coordinates an unanticipated Glu ligand contributed by an auxiliary helix implicated in substrate binding by docking and molecular dynamics simulation. The additional ligand is conserved in another experimentally validated HO-like *N*-oxygenase but not in two known HO-like diiron desaturases. Among ∼9600 sequences identified bioinformatically as belonging to the emerging HO-like diiron protein (HDO) superfamily, ∼25% have this carboxylate residue and are thus tentatively assigned as *N*-oxygenases.

**Significance statement:** The enzyme SznF assembles the *N*-nitrosourea pharmacophore of the drug streptozotocin. Its central *N*-oxygenase domain resembles heme-oxygenase (HO) and belongs to an emerging superfamily of HO-like diiron enzymes (HDOs) with unstable metallocofactors that have resisted structural characterization. We investigated assembly of the O_2_-reactive diiron complex from metal-free SznF and Fe(II) and leveraged this insight to obtain the first structure of a functionally assigned HDO with intact cofactor. Conformational changes accompanying cofactor acquisition explain its instability, and the observation of an unanticipated glutamate ligand that is conserved in only a subset of the HDO sequences provides a potential basis for top-level assignment of enzymatic function. Our results thus provide a roadmap for structural and functional characterization of novel HDOs.

---

Enzymes with histidine- and carboxylate-coordinated dinuclear iron cofactors catalyze biologically, biomedically, and environmentally important reactions in organisms from bacteria to humans (1, 2). Most of these enzymes use dioxygen as a co-substrate and cleave strong C–H, O–H, or N–H bonds in their reactions. Extensively studied members of this functional class share the conserved four-helix-bundle architecture initially recognized in the iron-storage protein, ferritin (Fig. S1) (3). Examples of ferritin-like diiron oxidases and oxygenases (FDOs) include the β subunit of class Ia ribonucleotide reductase (4, 5), soluble methane monooxygenase (sMMO) (6-10), stearoyl acyl carrier Δ^9^ desaturase (Δ^9^D) (11, 12), toluene/*o*-xylene monooxygenase (ToMO) (13-16), and arylamine *N*-oxygenases from the biosynthetic pathways that build the antibiotics aureothin (AurF) (17-20) and chloramphenicol (CmlI) (21-25). Each of these proteins reduces O_2_ at its Fe_2_(II/II) cofactor to form a *µ*-peroxo-Fe_2_(III/III) intermediate (1). In some cases (e.g., sMMO) additional proteins (MMOB) must bind to the FDO component (MMOH) to activate its cofactor for efficient O_2_ capture (8), and, in others (e.g., Δ^9^D) (11), binding of the substrate triggers this step. Early studies emphasized the conversion of the relatively inert initial peroxide-level intermediates to more reactive high-valent [Fe(IV)-containing] complexes by reductive or redox-neutral O–O bond cleavage (4, 5, 26-29), but later studies, including those on the *N*-oxygenases, suggested reaction of substrates directly with the initial mid-valent peroxide adducts (18-21, 23). In these cases, activating protonation and/or isomerization of the canonical *µ*-1,2-peroxide core was generally posited. Regardless of the pathway, the reactions of these proteins most commonly generate *µ*-(hydr)oxo-Fe_2_(III/III) product clusters, which must be reduced in situ by electron transfer proteins (e.g., ferredoxins or dedicated reductase components) for subsequent turnovers (1, 2). Thus, the diiron clusters of the FDOs act as stable cofactors, cycling through multiple redox states in discrete oxidative and reductive half-reactions.

In the last decade, O_2_-reactive, His/carboxylate-coordinated diiron cofactors have been identified in several other protein architectures, including the HD domain (30), the metallo-β-lactamase fold (31), the TIM-barrel fold (32, 33), and a HEAT-repeat domain (34, 35). Another new structural type, the seven-helix architecture first recognized in heme oxygenase (HO) (Fig. S1) (2, 36), has emerged recently as a second privileged diiron scaffold that may rival the ferritin-like fold in the versatility of transformations it supports. At least four HO-like diiron enzymes (HDOs) – a pair of *N*-oxygenases (37-40) and a pair of olefin-forming lyase/oxidase (hereafter, desaturase) enzymes (41-44) – have been described to date, and several others have been identified within biosynthetic gene clusters (Fig. S2) (45-48). The presumptive enzymatic activity of the first structurally characterized protein in this new superfamily, <u>C</u>hlamydia protein <u>a</u>ssociated with <u>d</u>eath <u>d</u>omains (CADD), is still unknown (49, 50). The first HDO to be assigned an enzymatic function, UndA, oxidatively decarboxylates dodecanoic (lauric) acid to undecene (41). An x-ray crystal structure of UndA revealed only a single iron ion with His_2_/carboxylate protein coordination within the relatively open HDO core, and it was initially suggested that UndA uses the observed monoiron(II) center for O_2_ activation (41). A pair of later studies showed, by contrast, that the iron cofactor is dinuclear (42, 43). One of these studies characterized a *µ*-peroxo-Fe_2_(III/III) complex on the reaction pathway (43). In that work, concerted efforts to solve a structure with both cofactor subsites fully occupied were unsuccessful, and regions of the protein implicated in providing ligands for the second iron site (core α3) gave weak electron density, indicative of structural disorder. In roughly concurrent studies, the aforementioned pair of HO-like *N*-oxygenases, involved in biosynthesis of streptozotocin (SznF) (37, 38) and azomycin (RohS) (40), also failed to retain their predicted diiron cofactors. For the case of SznF, multiple attempts at cofactor reconstitution, either prior or subsequent to crystallization, did not yield a high-quality structure of the holoenzyme, although partially metallated forms were successfully characterized (Fig. S3) (37). The apparent conservation of cofactor instability among the first several functionally assigned proteins in the emerging HDO superfamily (37, 38, 40, 41, 43) was taken as evidence that they might function by a novel *modus operandi*, perhaps utilizing iron more as a labile co-substrate than as a stably bound, in-situ-recycled cofactor (39). Regardless of the reason, no structure of a catalytically relevant (diferrous or diferric) holoenzyme HDO complex for a functionally assigned member of this emerging superfamily has yet been reported.

SznF has, in addition to its HDO central domain, an N-terminal dimerization domain and a C-terminal mono-iron cupin domain (Fig. S4). The HDO domain sequentially hydroxylates each of the two unmethylated guanidino nitrogen atoms of *N*^ω^-methyl-L-arginine (L-NMA), with *N*^δ^ being targeted first, before the cupin domain rearranges the triply modified L-Arg to *N*^δ^-hydroxy-*N*^ω^-methyl-*N*^ω^-nitroso-L-citrulline (Fig. S2), the proposed source of the *N*-nitrosourea pharmacophore of streptozotocin (37). The first SznF structures had a single iron ion in the cupin domain coordinated (in a mode potentially similar to that required for the final rearrangement step) by the *N*^δ^-hydroxy-*N*^ω^-methyl-L-arginine (L-HMA) product of the first HDO-mediated hydroxylation, but, as previously noted, did not reveal the structure of the HDO diiron cofactor (Fig. S3). In a later study, we showed that, far from being incompetent for iron binding, the HDO domain of the protein can readily acquire Fe(II) from solution and use it to capture O_2_ in a *µ*-peroxo-Fe_2_(III/III) complex (39). Acceleration of decay of this complex by mixing with either HDO substrate (L-NMA or L-HMA) showed that this complex either is the *N*-oxygenating complex for both steps or undergoes a substrate-promoted, rapid, reversible, and disfavored conversion to the oxygenating intermediate, which is then efficiently captured by the substrate. Here, we used the spectroscopic signature of this intermediate (absorption feature centered at 629 nm with molar absorptivity of ∼ 3 mM^−1^cm^−1^) to report on assembly of O_2_ reactive Fe_2_(II/II) complex from the apo protein and Fe(II) and kinetically dissect the assembly process. We then further leveraged this understanding to solve the structure of SznF with its diiron cofactor intact, thus providing the first look at a mechanistically relevant reactant complex of any member of this emerging enzyme superfamily. Comparison of an improved structure of SznF with its HDO domain in its metal-free (apo) form to that of the Fe_2_(II/II) complex affords a detailed structural description of the process of Fe(II) acquisition and the major protein conformational changes accompanying it. Together, the results provide a structural rationale for the cofactor instability seen in each of the initially described HDO systems.

## Results and Discussion

### Pathway to the µ-Peroxo-Fe_2_(III/III) Intermediate in SznF from Apo Protein, Fe(II), and O_2_

We recently demonstrated accumulation of a *µ*-peroxo-Fe_2_(III/III) intermediate to a high level (∼ 0.65 enzyme equivalents) upon exposure of a solution containing apo SznF and 3 equiv Fe(II) to excess O_2_ (39). The intermediate exhibits intense long-wavelength absorption (λ_max_ = 629 nm, ε_629_ ∼ 3 mM^−1^cm^−1^), which presumably arises from a peroxide-to-iron charge-transfer (LMCT) transition. In this study, we used development of this feature as the readout to dissect cofactor assembly and intermediate formation. By varying the concentrations, the molar ratios, and the order of mixing of the apo protein, Fe(II) and O_2_ reactants, we resolved four steps in this process (Scheme 1A). The apo protein binds Fe(II) tightly at one diiron subsite, and this first, tight-binding step requires a relatively slow (< 40 s^−1^ at 5 °C) protein conformational change [either before or after Fe(II) association] that partly rate-limits formation of the *µ*-peroxo-Fe_2_(III/III) intermediate from the apo protein and free Fe(II). The second iron subsite has modest affinity and is not saturated even with free Fe(II) concentrations in the 10^−4^–10^−3^-M range. Fe(II) binding at the less avid subsite is also relatively sluggish (∼1.5-3.0 × 10^3^ M^−1^•s^−1^) and is followed by more rapid capture of O_2_ (∼ 1.5-5 × 10^5^ M^−1^•s^−1^). The data also suggest that the associated monomers of the homodimer interact anti-cooperatively to engender partial “half-of-sites” reactivity (Scheme 1B), which manifests differently in initiation of the reaction by either (i) mixing of the pre-formed Fe(II)•SznF complex with O_2_ or (ii) simultaneous mixing of the apo protein with Fe(II) and O_2_. We summarize the data revealing this complex cofactor-assembly mechanism and likely anti-cooperativity in the following sections before taking up the structural basis for the effects.

**Scheme 1.**
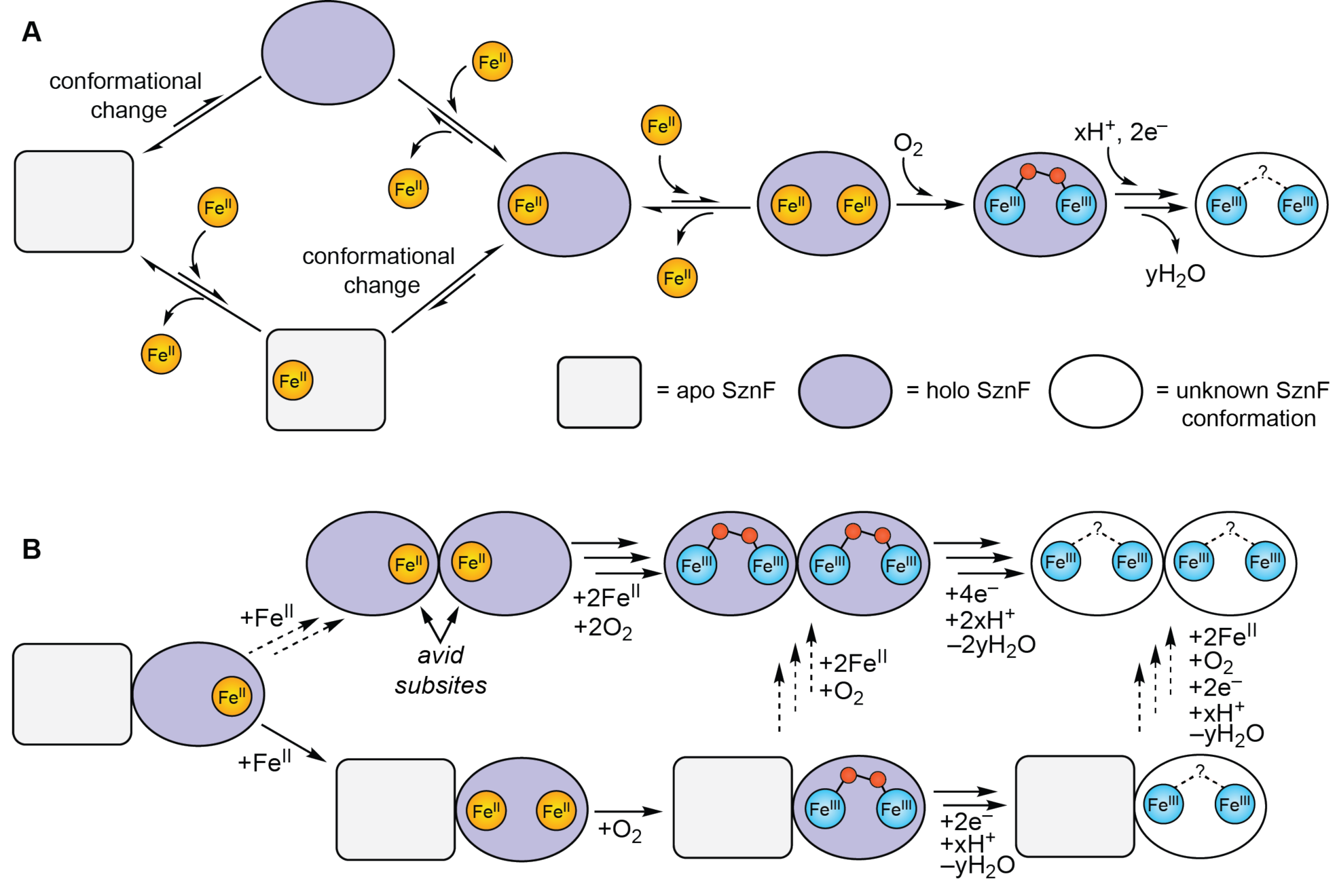
Assembly and reaction of the SznF diiron cofactor. (*A*) A conformational change accompanies Fe(II) binding to the avid subsite in the HDO domain of SznF. (*B*) Rationale for the divergent kinetic behavior of reactions initiated from pre-formed Fe(II)-SznF complexes (*top pathway*) and the apo protein (*bottom pathway*).

### Effect of Fe(II):Protein Ratio on the Kinetics of the Reaction of the Fe(II)•SznF(-Δcupin) Complex with O_2_

The kinetics of formation and decay of the *µ*-peroxo-Fe_2_(III/III) intermediate in reaction of the pre-formed Fe(II)•SznF complex with O_2_ depend in a complex way on the metal-to-protein ratio in the anoxic reactant solution (Fig. 1A). Several informative trends can be seen in these stopped-flow absorption traces. The reaction with Fe(II):SznF = 1 fails to accumulate the *µ*-peroxo-Fe_2_(III/III) intermediate (Fig. 1A, *blue*). Accumulation of the intermediate increases with Fe(II):SznF ratio in the range of 2-4 (Fig. 1A, *red, orange, green, cyan, purple*), and “saturation” sets in as the ratio reaches 6.0 (Fig. 1A, *black*). The data imply that the O_2_-reactive dinuclear complex is not populated when the apo protein is allowed to reach equilibrium with equimolar Fe(II). Mixing of the anoxic Fe(II)•SznF reactant solution with an anoxic solution of the colorimetric Fe(II) indicator ferrozine (λ_max_ = 562 nm, ε_562_ = 27.9 mM^−1^cm^−1^ (51, 52), rather than with O_2_-saturated buffer, revealed that, with Fe(II):SznF = 1, essentially all the Fe(II) is bound (Fig. S5). Free Fe(II) is chelated rapidly (*k*_obs_ > 50 s^−1^) by ferrozine under the conditions of this experiment, engendering a fast phase of development in the *A*_562_ kinetic trace. There is no such rapid phase with an Fe(II):SznF ratio of 1.0 (Fig. S5A-B, *blue*), but increased ratios give steadily increasing amplitude attributable to free Fe(II) (Fig. S5A-B, *red, green, cyan, purple, black*). In principle, tight binding in the cupin domain of the wild-type enzyme could fully sequester one equiv Fe(II) to prevent its binding in the HDO domain, thus preventing formation of the peroxo-Fe_2_(III/III) complex upon exposure of the Fe(II)•SznF reactant to O_2_. However, the variant protein with all three His ligands (407/409/448) in the cupin domain replaced by Ala (SznF-Δcupin) behaves similarly, exhibiting neither accumulation of the *µ*-peroxo-Fe_2_(III/III) complex (upon mixing with O_2_) nor a fast phase of chelation by ferrozine when the anoxic protein reactant solution contains equimolar Fe(II) and protein (Fig. 1B and Fig. S5C-D). It thus appears that binding of Fe(II) in the cupin domain does not contribute to the observed complex behavior, and we hereafter denote Fe(II)•SznF complexes in terms of only the two sites contributed by the HDO domain. These observations suggest that pre-incubation of apo SznF(-Δcupin) with one equiv Fe(II) leads to sequestration of the metal in an unreactive mononuclear complex *with the metal bound in one of the two cofactor subsites of the HDO domain*. This situation would arise, most simply, from substantially different intrinsic affinities of the two subsites and only modest (or no) cooperativity between them.

**Fig. 1.**
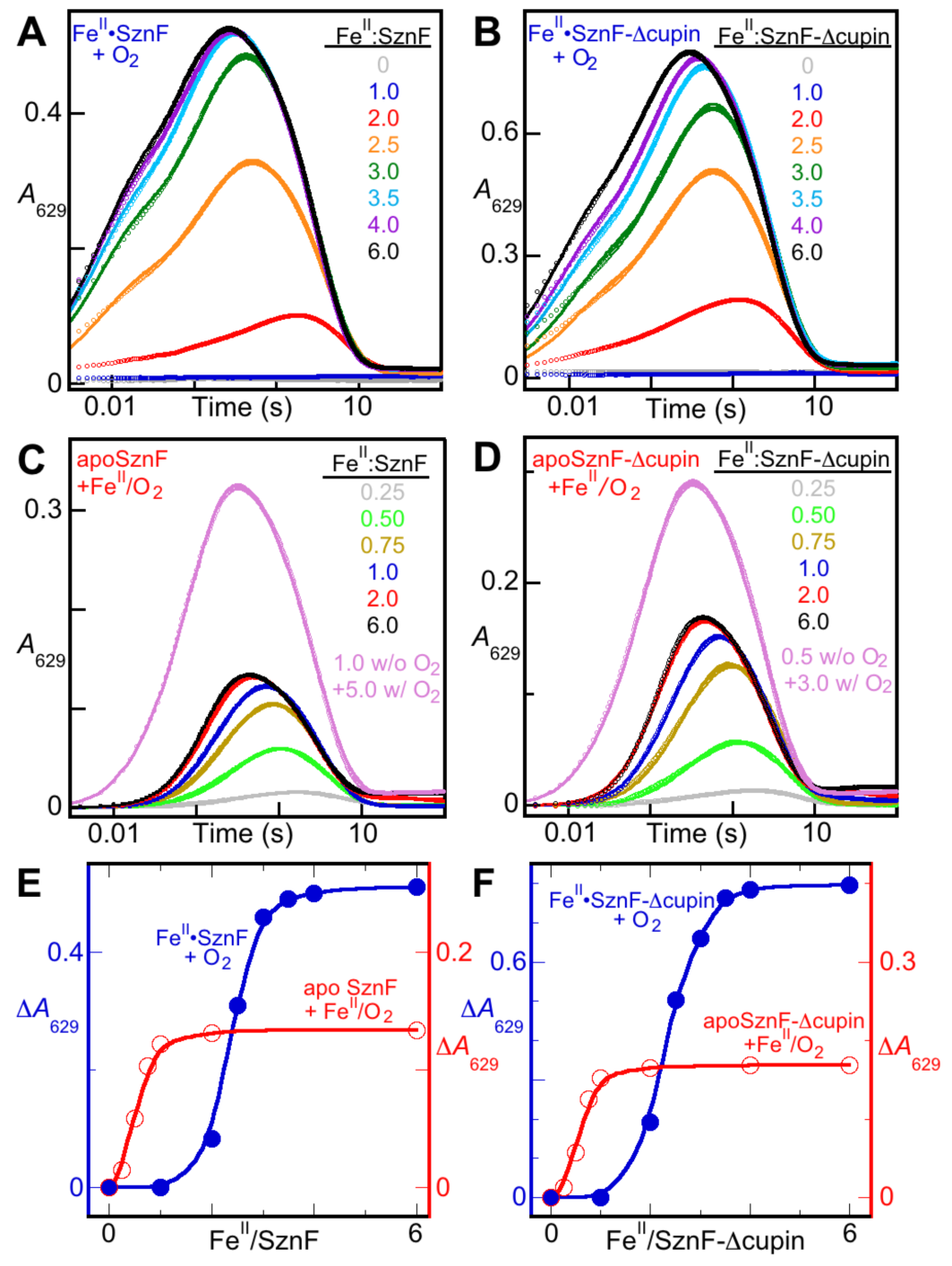
Stopped-flow absorption experiments monitoring assembly of the O_2_-reactive Fe_2_(II/II) cofactor in the HDO domain of SznF(-Δcupin) by its conversion to the intensely absorbing *µ*-peroxo-Fe_2_(III/III) intermediate. (*A*) Absorbance at 629 nm (*A*_629nm_) versus time after an anoxic solution containing wild-type SznF and Fe(II) was mixed at 5 °C with an equal volume of O_2_-saturated buffer. (*B*) Traces from corresponding reactions of the Δcupin variant. The solid lines in *A* and *B* are fits of the equation (see *SI Appendix*) appropriate for the model of Scheme S1 to the data. (*C, D*) Traces from reactions of a solution of SznF(-Δcupin) containing 0-1 equiv Fe(II) with (additional) Fe(II) and O_2_. The solid lines are fits of equation 1 (*SI Appendix*) to the data. (*E, F*) Differences between maximum and minimum *A*_629nm_ for the traces in *A*-*D* plotted versus Fe(II):SznF(-Δcupin). The solid lines are interpolation fits to the data. When no Fe(II) was present in the protein solution, it was saturated with O_2_, and the Fe(II) solution was anoxic and in buffer. When Fe(II) was present in the protein solution, it was anoxic and the Fe(II) solution was both O_2_-saturated and in 1 mM H_2_SO_4_ rather than buffer.

A second insight that is obvious from inspection of the Δ*A*_629_ traces, particularly those from the reactions with Fe(II):SznF = 2.5 – 6, is that formation of the intermediate is kinetically biphasic. The faster phase of development, complete within ∼ 0.04 s, acquires increasing amplitude with increasing Fe(II):SznF, coming to represent ∼ half of the total rise phase at the highest Fe(II):SznF ratio examined (Fig. S6A). The apparent first-order rate constant associated with this phase, extracted by fitting equation 1 (appropriate for Scheme S1; see below) to the data (solid lines in Fig. 1A), varies by less than 10% from the mean of 1.5 × 10^2^ s^−1^ for all Fe(II):SznF ratios tested from 2.5 to 6 (Fig. S6B). In this range, free Fe(II) – reflected by the fast phase of chelation in the anoxic mix with ferrozine – increases proportionally to the Fe(II):protein ratio (Fig. S5B, triangles). The kinetic insensitivity of the fast phase of intermediate formation to the concentration of free Fe(II) suggests that it is associated with trapping of O_2_ by the dinuclear Fe_2_(II/II)•SznF complex already formed in the anoxic pre-incubation. The slower phase in the *A*_629_-versus-time traces becomes faster and loses amplitude as the Fe(II):SznF ratio increases from 3 to 6 (Fig. S6A). Its apparent rate constant from the fitting analysis (*k*_1_’) nearly doubles (from 6 to 12 s^−1^) across this range (Fig. S6B, triangles), implying that intermediate formation in the slow phase requires a bimolecular step involving free Fe(II). Succinctly, the more rapid phase of intermediate formation reflects protein with its O_2_-reactive Fe_2_(II/II) cofactor fully assembled, and the slower phase reflects protein with only the more avid subsite occupied (Scheme 1). The fraction of the protein in the mononuclear form decreases as Fe(II):SznF increases (because the less avid subsite is increasingly occupied), and the amplitude associated with the slower, [Fe(II)]-dependent phase of intermediate formation thus decreases. The other effect of increasing Fe(II):SznF is that any mononuclear complex that still remains can proceed more rapidly through the two-step sequence of (i) Fe(II) binding at the less avid subsite and (ii) O_2_ addition, because the concentration of free Fe(II) rises proportionally (as shown by the ferrozine experiment) and the effective first-order rate constant for association at the less avid site increases accordingly.

### Kinetics of the Reaction of Oxygenated Apo SznF(-Δcupin) with Fe(II)

The kinetic behavior of the reaction initiated by mixing oxygenated apo SznF(-Δcupin) with Fe(II) is drastically different from that seen upon initiation from the pre-formed complex (Fig. 1C, F). First, there is a pronounced lag phase in formation of the intermediate, regardless of the Fe(II):protein ratio. The lag phase is not seen in the reaction of the pre-formed complex with O_2_ which implies that it is associated with a slow step in acquisition of Fe(II) by the apo protein. The fact that, even at very low Fe(II):SznF(-Δcupin) ratios (0-1), which do not support accumulation of intermediate in reaction of the pre-formed complex with O_2_, give considerable intermediate accumulation when starting from apo protein implies that the slow step is associated with sequestration of Fe(II) in the tight site. In other words, with time given for this slow step to reach completion in the absence of O_2_, equimolar Fe(II) ends up sequestered in only the more avid subsite, but, with O_2_ present from the start, filling of the avid subsite is followed by a two-step sequence of Fe(II) acquisition by the weaker site and addition of O_2_, and this sequence is faster than the initial binding in the avid subsite. One outcome of this situation is an efficient use of even sub-stoichiometric Fe(II) for still relatively rapid (despite the lag phase) intermediate formation when apo protein is mixed with Fe(II). A second, remarkable outcome is that an Fe(II):SznF(-Δcupin) ratio of 2 is essentially “saturating” in mixing apo protein with Fe(II) and O_2_; an increase to a ratio of 6 results in a very modest effect on the kinetics of the intermediate, in marked contrast to the behavior of the reaction of the pre-formed complex with O_2_. Even a mere 1 equiv Fe(II), which supports essentially no intermediate accumulation if allowed to bind to the protein in advance of O_2_ exposure, supports accumulation of > 90 % of the maximum quantity possible if it is mixed with apo protein in the presence of O_2_. In essence, “saturation” – as reported by the kinetics of the intermediate – requires much less Fe(II) if it is not allowed to reach equilibrium in advance of O_2_ exposure.

A second notable and (as yet) incompletely understood characteristic of the reaction as initiated from the apo protein is a markedly diminished amplitude in the *A*_629_-versus-time traces. Regression analysis of the traces according to Eq. 1 returned rate constants of ∼ 40 s^−1^ for the lag phase and 2-8 s^−1^ [varying with Fe(II):SznF(-Δcupin)] for the formation phase, with a maximum amplitude of 0.15 (0.19 for SznF-Δcupin). This amplitude is only ∼ one-fourth the value seen in the traces of Fig. 1A-B. Put simply, two steps occurring with these best-fit rate constants would not limit accumulation to the drastic extent seen in this comparison. A potential explanation lies in the dimeric nature of SznF and the precedent from the ferritin-like diiron enzymes of “half-of-sites reactivity.” Prior filling of the more avid subsites in both paired monomers of the dimer may allow for both monomers to form the intermediate rapidly (in either the fast or slow phase seen in Fig. 1A-B), whereas the cascade of Fe(II) binding at the less avid subsite and O_2_ addition to the dinuclear cluster that ensues after the apo protein dimer first binds Fe(II) in the more avid subsite may, by conformational coupling, block reaction of the other, paired monomer. In principle, the two phases of intermediate formation seen in reaction of the pre-formed Fe(II)•SznF(-Δcupin) complex with O_2_ could similarly represent reactions of pair monomers, the first having both subsites filled and the second requiring Fe(II) binding to the less avid subsite prior to O_2_ addition. In this scenario, conformational coupling across the dimer could engender a binding anti-cooperativity that serves to suppress formation of fully loaded Fe(II)_4_•SnzF dimer (Scheme S1A), as seen before in the ferritin-like class Ia ribonucleotide reductase β subunit from *E*. *coli* (53, 54). The fact that the amplitude for the Δcupin variant is greater than that for the wild-type protein may also relate to a perturbation of intermonomer coupling caused by replacement of all three His ligands with Ala, although an increased molar absorptivity of the intermediate in the variant scaffold is also theoretically possible.

### A Rate-Limiting Conformational Change Associated with Fe(II) Binding in the Avid Subsite

The results of a hybrid experiment, in which SznF(-Δcupin) was first incubated with ≤ 1 equiv Fe(II) in the absence of O_2_ and the resultant complex was then mixed simultaneously with additional Fe(II) and O_2_, confirm that binding in the avid subsite drives or locks in a conformational change that is responsible for the lag phase in intermediate formation in the reaction of apo protein with Fe(II) and O_2_. The quantity of Fe(II) in the anoxic pre-incubation in this experiment does not, by itself, support intermediate accumulation upon subsequent exposure to O_2_ (Fig. 1A-B, *blue*, and Fig. S7). Furthermore, the larger quantity of Fe(II) mixed (along with O_2_) with the resultant complex would, if added directly to the apo protein, support only the drastically diminished accumulation noted previously (Fig. 1C-D, *black*, and Fig. S7). It is therefore informative that addition of Fe(II) in *both* stages removes the pronounced lag in intermediate formation and allows ∼ twice as much to accumulate as in the reaction of apo protein directly with the same quantity of Fe(II) (Fig. 1 C,F, *pink*, and Fig. S7). Prior binding just to the avid subsite in the absence of O_2_ accomplishes the conformational change in advance of mixing with additional Fe(II) and O_2_, while leaving the second, less avid subsite still unoccupied. Mixing with Fe(II) and O_2_ then allows intermediate formation to proceed with the effective first-order rate constant characteristic of Fe(II) binding to the less avid subsite (∼12 s^−1^) and only a much shorter lag phase attributable to fast O_2_ addition (1.8 – 4.5 x 10^2^ s^−1^ with ∼ 0.9 mM O_2_, Fig. S8).

### X-ray Crystal Structure of SznF with an Intact Diiron Cofactor Reveals a Seventh Cofactor Ligand in the HDO Domain

In previously reported structures of SznF (PDB accession codes 6M9S, 6M9R), the HDO domain was largely devoid of iron (Fig. S3), despite concerted efforts to isolate the protein with an intact oxidized cofactor or reconstitute with Fe(II) and oxygen (37). Our recognition in a recent study that the Fe_2_(III/III) complex is unstable (39), and our elucidation here of the complex Fe_2_(II/II) cofactor assembly dynamics, suggested that prior crystallization of the apo protein followed by loading of Fe(II) *in crystallo* might allow for structural characterization of the holoenzyme complex. As recently reported, we produced the protein in *E*. *coli* cultures supplemented with Mn(II) to limit Fe(II) uptake during over-expression and purification and chelated the Mn(II) to produce the naive [not previously reacted with Fe(II) and O_2_] apo protein. We then crystallized the protein anoxically and briefly soaked crystals in a mildly acidic solution of Fe(II) for 5-10 min before freezing them for collection of x-ray diffraction datasets (*SI Appendix*, Table S1). This approach yielded a 1.66-Å resolution crystallographic model with high occupancy (67-100%) of all four Fe(II) binding sites in the two HDO domains of the SznF dimer. The identity of the Fe_2_(II/II) cofactor was validated by collection of anomalous diffraction datasets at the Fe and Mn x-ray *K*-edge absorption energy peaks. (Fig. 2A and Fig. S4). Strong peaks in the anomalous difference map were observed only in the datasets acquired at the Fe absorption energy. The structure confirms that the diiron cofactor is held within core helices 1-3 by the conserved 3-His/3-carboxylate ligand set (Fig. S1) predicted from alignment of its sequence with those of other characterized HDOs (Fig. 3).

**Fig. 2.**
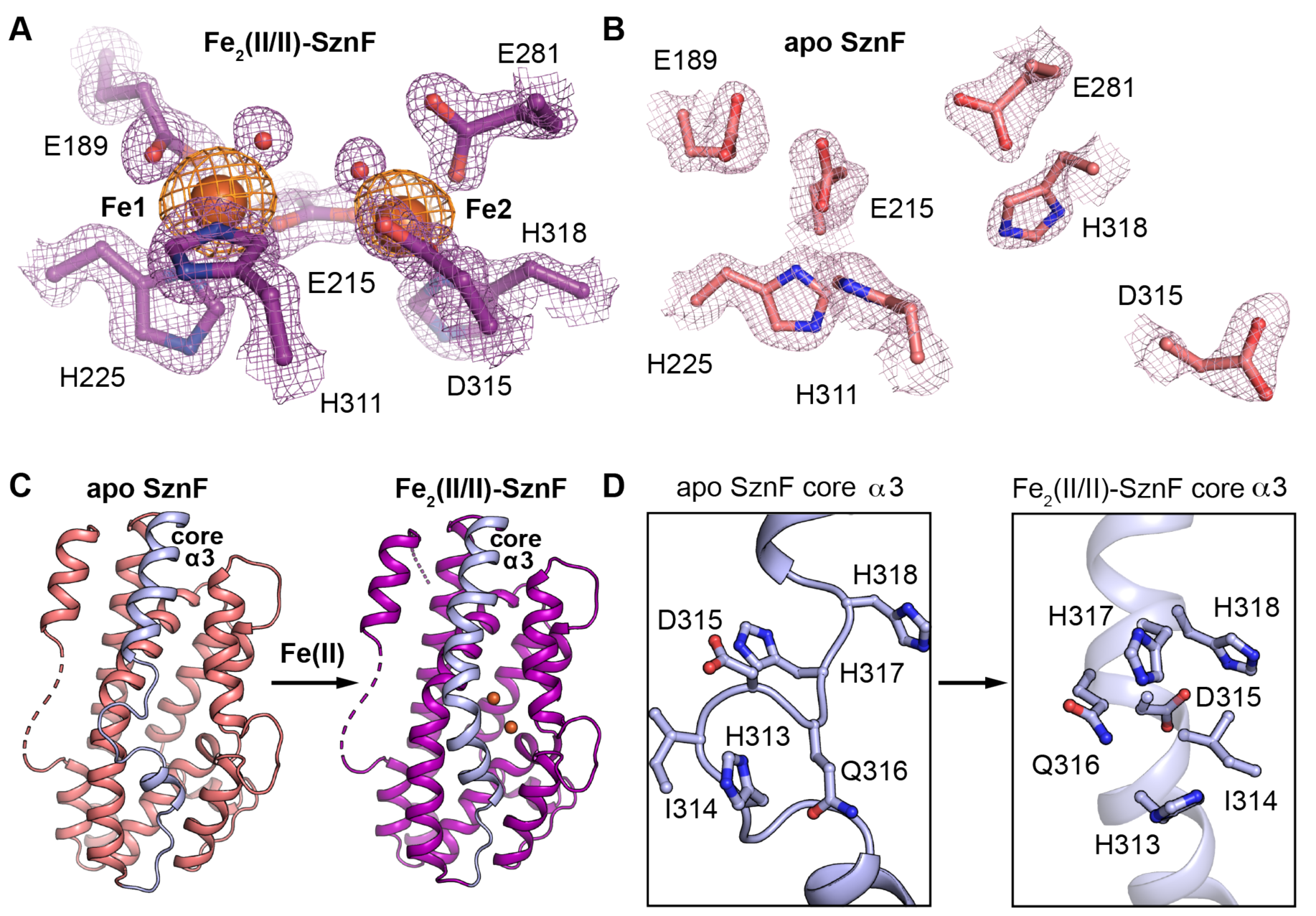
X-ray crystal structures of Fe_2_(II/II)-SznF (*A*) and apo SznF (*B*) reveal a conformational change in core α3 (*C, D*) upon metal binding. 2*F*_O_-*F*_C_ maps (purple or pink mesh) are shown contoured at 1.5σ (Fe_2_(II/II)-SznF) or 1.0σ (apo SznF). An iron anomalous difference map (orange mesh) is shown contoured at 3.0σ. Selected amino acid side chains are shown in stick format. Fe(II) ions and water molecules are shown as orange and red spheres, respectively.

**Fig. 3.**
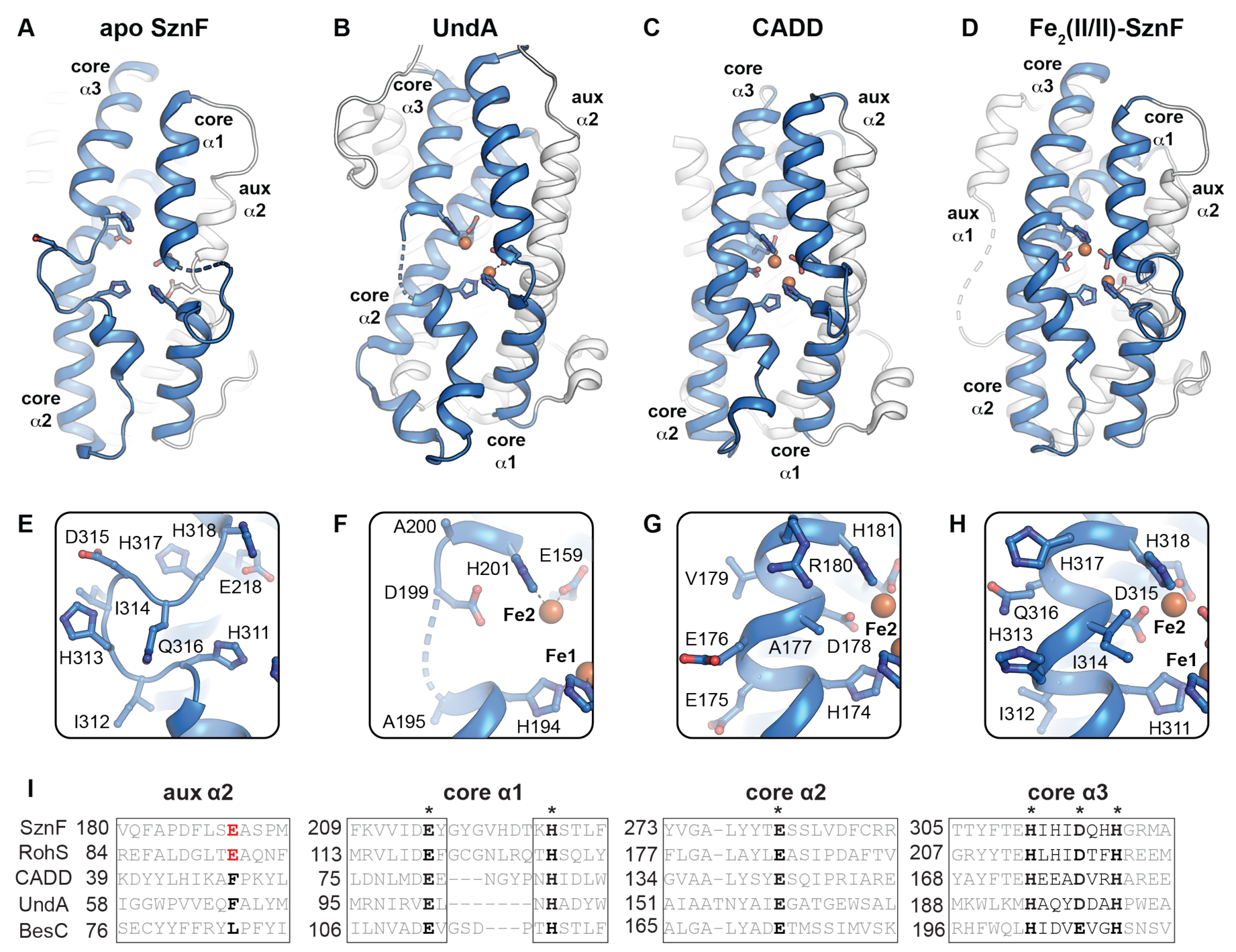
Structural comparison and sequence alignment of functionally diverse HDO proteins. Apo SznF (*A, E*) (PDB accession code 6XCV) and UndA (*B, F*) (PDB accession code 6P5Q) x-ray structures lack full-occupancy cofactors and exhibit disrupted or disordered regions in core α3. CADD (*C, G*) (PDB accession code 1RCW) and the iron-bound form of SznF (D,H) (PDB accession code 6VZY) contain full-occupancy diiron cofactors and maintain a canonical α-helix structure throughout core α3. A sequence alignment (*I*) of characterized *N*-oxygenases (SznF, RohS), oxidative desaturases (BesC, UndA), and protein of unknown function (CADD) shows that the newly identified Glu ligand in aux α2 of SznF is conserved in the *N*-oxygenases but absent in CADD and the desaturases. However, all enzymes conserve branched and polar amino acids in the metal-binding region of core α3, suggesting that all HDOs characterized to date are equipped to employ the conformationally-gated metal-binding mechanism described here for SznF.

In addition to verifying the six that were predicted, the new structure identifies an additional iron ligand, Glu189. CADD and UndA – two other structurally characterized HDOs – lack a coordinating residue at this position (Fig. S9). The SznF Glu189 carboxylate binds in bidentate mode to Fe1 (Fig. 2A and Fig. S10). Interestingly, FDO *N*-oxygenases, such as AurF and CmlI, have an analogous extra Fe1 ligand relative to their C/O-H-cleaving counterparts (Figs. S1, S11) (17, 22). However, the identity and origin of the additional ligand is different in the two scaffolds. In the FDOs, the extra ligand is a His provided by the same core helix (core α4) that harbors the Glu and His ligands in the second of the tandem pair of EXXH motifs that identifies the scaffold (Fig. S1). In SznF, Glu189 is instead contributed by one of the auxiliary helices (aux α2) found in the HDO scaffold (Figs. S1, S11).. Despite these differences, the extra ligand serves to differentiate the cofactor subsites in both scaffolds, giving a higher coordination number for Fe1 than for Fe2. It has been suggested for AurF and CmlI that saturation of the Fe1 coordination sphere by the extra His disfavors the η^2^ interaction of the dioxygen unit with this iron that would be required for O–O-bond cleavage to generate a high-valent state, thus stabilizing the peroxide-level complex for its direct oxidization of the substrate nitrogen (22). This hypothesis seems reasonable also for the case of SznF.

In the Fe_2_(II/II) SznF cofactor, the six-coordinate Fe1 has a distorted-octahedral geometry, whereas the five-coordinate Fe2 has roughly square-pyramidal symmetry. Each site additionally coordinates a water molecule trans to a His ligand (Fig. 2A, Fig. S10). Interestingly, these presumably labile waters are separated by a distance of only 2.8 Å, and they project toward one another into a solvent-lined cavity that could potentially harbor the L-NMA or L-HMA substrate (see below). As such, their locations might approximate the positions of the O-atoms of the bridging peroxide in the detected intermediate. However, formation of the *µ*-peroxo-Fe_2_(III/III) intermediate would likely require significant contraction along the Fe–Fe vector: the ∼ 5.0 Å separation observed in the structure is much greater than those found in most structurally characterized *µ*-peroxo-Fe_2_(III/III) complexes with similar ligand sets (2.5 – 3.4 Å) (1). This long intermetallic separation in reduced SznF is enforced by both local and global structural features. The cluster has a single bridging ligand, the carboxylate of Glu215 from core α2, which coordinates in a *µ*-1,3 mode. By contrast, reduced complexes of FDOs generally have two or three bridging carboxylates and intermetallic distances of less than 4 Å (1). In some cases, a μ-η^1^,η^2^ mode of one of these carboxylates is associated with contraction along the Fe–Fe axis, and this geometry is proposed to be diagnostic of the O_2_-reactive state (55). No such bridge is present in reduced SznF (Fig. S11).

The long Fe–Fe vector in SznF may also arise, in part, from irregularities within the helices that provide the ligands. Of the four helices that provide ligands, two – aux α2 and core α1 – contain significant loop/turn interruptions, and the other two – core α2 and core α3 – exhibit more subtle deviations from canonical backbone H-bonding that result in curvature of the helical axis. These irregularities would, in general, be expected to increase flexibility in the vicinity of the cofactor (Fig. S12). Interestingly, 3 of the 4 ligand-providing helices of the HDO scaffold face the solvent. Their surface exposure, coupled with the aforementioned structural distortions, likely underpins the unusual iron-binding properties of SznF. Furthermore, the large separation of the Fe(II) ions and single carboxylate bridge connecting them could explain why the two cluster subsites behave essentially as one tighter (cofactor-like) subsite and one weaker (substrate-like) subsite, with no obvious cooperativity between them.

### Visualization of a Conformational Change in Core Helix 3 Accompanying Cofactor Assembly

Comparison of the occupancies of the two cofactor subsites – 90-100% for subsite 1 versus 60-80% for subsite 2 – implies that subsite 1 has the greater affinity. This assignment, which would appear to correlate with the greater coordination number of Fe1, would implicate Fe(II) binding to subsite 1 in driving the conformational change. To elucidate the nature of this change, we also solved the structure of the naive apo protein, omitting the Fe(II) soak before freezing the crystals for data collection. The resulting diffraction datasets, collected to a resolution of 1.8 Å, yielded more well-defined electron density for the protein than did the datasets collected in the prior study (Fig. 2B). Most notably, the entire segment spanning core helix α3 – both backbone and side chains – could be definitively modeled, whereas it had previously exhibited partial disorder (Fig. S3). In the new apo structure, the segment spanning residues 311-318, with three cofactor ligands (His311, Asp315 and His318), adopts an ordered, non-helical conformation. By contrast, in the iron-bound structure, this same segment is essentially α-helical (although three canonical backbone interactions are absent), and core α3 is uninterrupted. Remodeling of secondary structure in the residues spanning core α3 thus appears to be a key part of the conformational change seen in the kinetics of cofactor assembly in SznF (Fig. 2C, D).

In addition to this α3 backbone remodeling, multiple, substantial movements of cofactor-coordinating side chains are associated with iron binding. The histidine and carboxylate ligands contributed by aux α2, core α1, and α2 rotate. The ligands in core α3 change position even more drastically. Asp315 undergoes the greatest shift, moving ∼ 7 Å from the surface of the protein and folding into the cavity that harbors the cofactor. Interestingly, this side chain is a ligand only to Fe2, in the subsite of lesser affinity. Repositioning of Asp315 is likely a crucial part of the conformational change, as it should properly configure the second subsite. The data paint a detailed picture of the cofactor assembly process, in which loading of subsite 1 initiates refolding of core α3 to bring subsite 2 ligand Asp315 into position to coordinate the second Fe(II).

### Role of Core α3 Dynamics in Cofactor Instability of HDOs

Comparison of the structure of core α3 in the HDO domain of SznF to those of its counterparts in UndA and CADD (Fig. 3A-H) reveals correlations that may be functionally significant. CADD appears to be an outlier in the emerging superfamily (at least at this early stage) in having a stable cluster that allows it to be isolated with bound metal (49). As in Fe_2_(II/II)-SznF, core α3 of CADD, which we presume to have been characterized in its oxidized Fe_2_(III/III) state, is fully α-helical in the published structure. By contrast, UndA shares with SznF the property of cofactor instability and still has not been crystallographically characterized with its diiron cluster intact, despite concerted efforts toward this goal (43). Core α3 in the UndA structure is partially disordered, as in the previously reported structures of SznF (37), and the Asp315 equivalent, Asp198, resides in or near this unmodeled stretch (Fig. S9). However, the fact that substitution of this residue abolished activity in UndA was taken to imply that it is a cofactor ligand, which would, in turn, suggest that it must also be brought into position during assembly of the O_2_-reactive complex (43). Here, an additional event, fatty acid (substrate) binding, is necessary to produce the reactive state. In UndA, it is not known whether substrate binding precedes, interrupts, or follows binding of the two Fe(II) ions. It seems likely that metal binding in subsite 2 with coordination of the mobile core α3 carboxylate – brought into place by a conformational change driven by binding of Fe(II) (to subsite 1) and/or substrate – could be the final step, as in SznF. Moreover, the structural features associated with weak Fe(II) binding in subsite 2 may be conserved across the larger superfamily. Core α3 is completely surface exposed in all three structurally characterized HDOs. Analysis of the conformationally mobile segment in SznF core α3 (HIHIDQHH) shows that it contains entirely polar or β-branched side chains. Burial of such a sequence in the interior of an α-helix might be disfavored in the absence of compensating metal-ligand interactions. Alignment of the corresponding segments of the functionally assigned HDOs (Fig. 3I) shows that all are rich in polar or branched (or both) amino acids.

### Identification of the Substrate Binding Site in the SznF HDO Domain by Computational Modeling Guided by Structural Comparisons

Because initial efforts to obtain an experimental structure of SznF in complex with one of its substrates (L-NMA or L-HMA) were unsuccessful, we used computational docking and molecular dynamics to assess how L-NMA might bind and, importantly, how the regiochemistry of hydroxylation might switch. Cavity-mapping analysis (56) with the coordinates of the Fe_2_(II/II)-SznF structure revealed a pocket near the cofactor with sufficient volume (>1000 Å^3^) to accommodate the substrate (Fig. S13). The pocket resides on the same face of the metal cluster as the solvent ligands postulated to approximate the positions of the peroxide oxygen atoms in the intermediate. Substrate bound within this pocket would be ideally poised for direct nucleophilic attack on the peroxide, one mechanism proposed for the FDO *N*-oxygenases (18, 21). The SznF cavity is directly connected to the protein surface, terminating near a disordered six-residue sequence in aux α1. This opening provides a plausible access route for substrate to enter the HDO active site. FDO *N*-oxygenases are also purported to undergo a closed-to-open transition to gate substrate access to the active site, although the proposed access route is distinct (22).

Despite the ionic character of both SznF HDO substrates, this hypothetical binding site is largely hydrophobic. Only a single charged side chain, that of Asp185, projects into the pocket. As noted, it resides on aux α2, which also contains the newly identified Glu189 ligand. Aux α2 is interrupted near these side chains by a type II β-turn insertion (Fig. 4A). Glu189 is located at the tip of this β-turn. The motif both enables a closer approach of the coordinating carboxylates and exposes additional backbone polar functional groups that could interact with the substrate. This interruption could at least partly complement the charges of the substrates via helical dipoles. Specifically, the bisection of aux α2 by the β-turn exposes negative and positive dipoles that could orient near the α-ammonium and α-carboxylate groups, respectively. The turn is flanked at its C-terminus by Met193. Its thioether side chain projects into the active site. Along with other hydrophobic groups in the substrate pocket, the Met193 side chain could contribute to selectivity for the *N*-methylated L-Arg over the more prevalent, unmethylated form. Intriguingly, the next residue, Met194, projects away from the active site toward its counterpart in the opposite monomer, where it could play a role in the proposed cross-dimer anti-cooperativity in Fe(II) binding (Fig. S14). Thus, aux α2 not only delivers a ligand that was not anticipated from sequence comparisons with other HDOs but may also (i) coordinate intermonomer communication relevant to cofactor assembly and (ii) provide multiple substrate binding determinants.

**Fig. 4.**
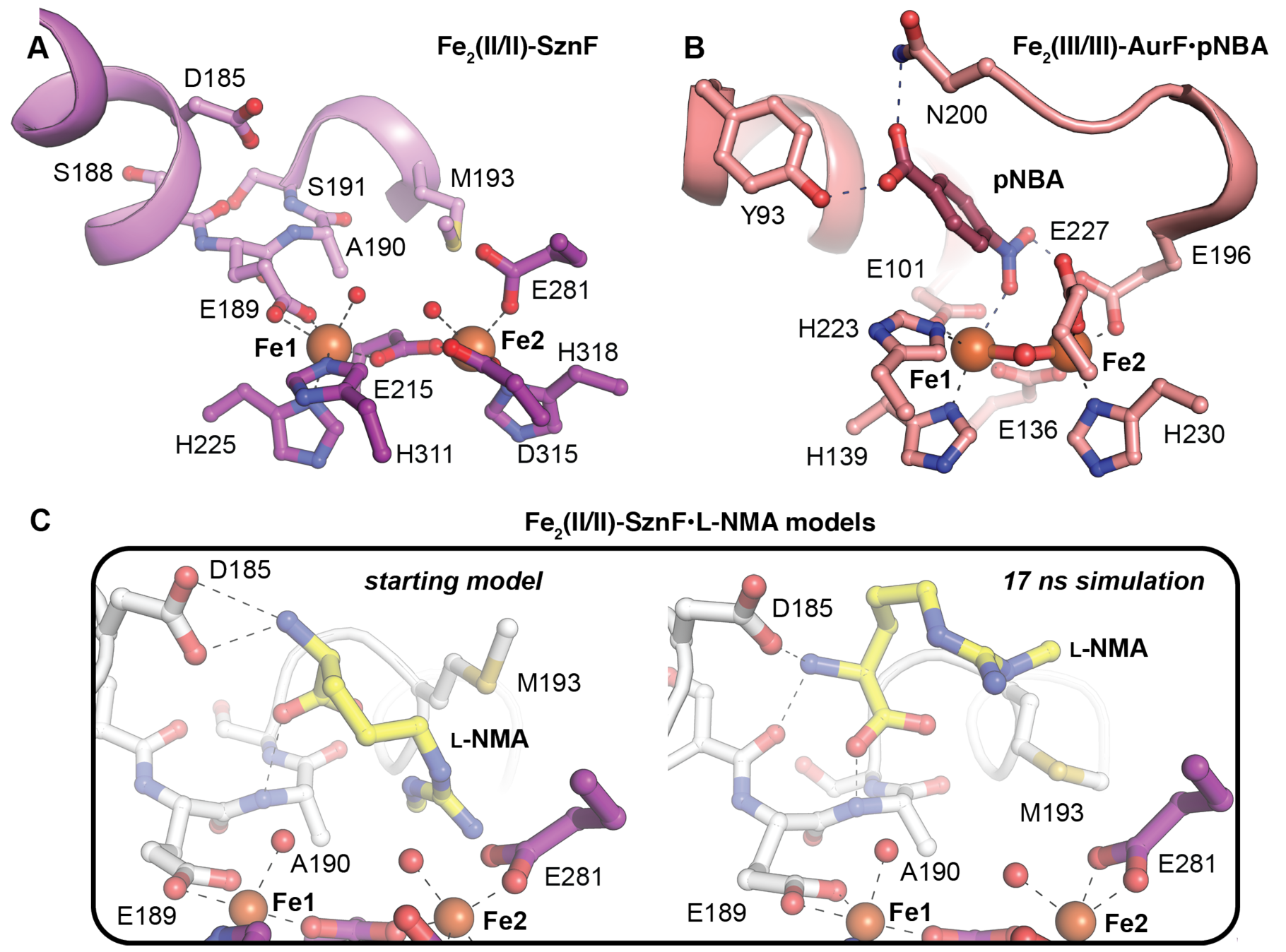
Aux α2 (violet in panel *A*, white in panel *C*) contributes an unanticipated seventh ligand and predicted substrate-binding determinants. A comparison of the metal coordination environment and putative substrate or product binding pockets in SznF (PDB accession code 6VZY) (*A*) and AurF (PDB accession code 3CHT) (*B*) reveals similarities between the HDO and FDO *N*-oxygenases. An energy minimized docking model of the SznF substrate, L-NMA, before (*left view*) and after (*right view*) molecular dynamics simulation (*C*) shows that predicted substrate H-bonding interactions in aux α2 can be accommodated. Selected amino acids and substrates are shown in stick format. Fe^II^ ions and water molecules are shown as orange and red spheres, respectively.

To test the idea that the aux α2 functional groups contribute to binding and proper orientation of L-NMA, we generated a docking model of the Fe_2_(II/II)-SznF•L-NMA complex. We reasoned that the α-amine and α-carboxylate groups might interact with the side chain of Asp185 and the backbone amide of Ala190, respectively. We subjected an energy-minimized docking model of this complex to molecular dynamics (MD) simulations (Fig. 4C and Fig. S15). The simulations show that the substrate can be accommodated in the open cavity above the cofactor while maintaining the predicted contacts with the Asp185 carboxylate and Ala190 amide nitrogen within the β-turn. While these interactions persist throughout the simulation, the L-NMA side chain remains dynamic, populating different rotamers and occupying different positions within the active site (Fig. S15). In general, the pair of guanidino nitrogen atoms that are hydroxylated in the sequential steps remain close to the cofactor; their relatively high mobility may enable the switch in regiochemistry (37, 38).

The modeled structure provides a basis to compare the location and nature of the substrate-binding pocket in SznF with those in other HDOs and FDOs. Some published structures of UndA include bound lauric acid (Figs. S9 and S16) (41, 43) and, although none of these have high occupancies of iron subsite 2, they do reveal direct coordination of the substrate carboxylate by Fe1, in approximately the same location as the Glu189 ligand in SznF. Correspondingly, aux α2 of UndA lacks the β-turn insertion that harbors the extra carboxylate ligand and is instead uninterrupted in all published structures (41, 43). The carboxylate of Glu189 in SznF and the substrate carboxylate in UndA might thus be, in a sense, functionally analogous. If this carboxylate-Fe1 interaction is necessary to set the geometry and tune the electronic structure of the cofactor for O_2_ capture, the difference between the UndA and SznF structures would explain why the O_2_-addition step is markedly accelerated by substrate binding in the former HDO (43) but not in the latter (39). Despite this difference, aux α2 of UndA, like the corresponding helix in SznF, contributes groups that interact with the substrate: the hydrophobic side chains of Val65 and Phe68 both contact the alkyl chain of the bound lauric acid (41, 43). The comparison suggests that analysis of aux α2 sequences in HDOs might aid in identifying their substrates and perhaps engineering their specificities and outcomes.

Although the FDO and HDO scaffolds have completely different topologies, the *N*-oxygenases in the two superfamilies appear to use a conserved coordination sphere (3-His/4-carboxylate) that, in both cases, includes an increased coordination number of their Fe1 site in comparison to their C/O-H activating counterparts (Figs. S1, S11, S16). However, in the FDOs, the extra ligand resides on a core helix that, across the entire superfamily, contributes two other cofactor ligands (17), whereas, in SznF, it extends from an auxiliary helix that contributes only this ligand, and only in a subset of the superfamily (Fig. S1). AurF is the only FDO *N*-oxygenase crystallographically characterized to date with its product bound (17). The two systems share a similar relative disposition of the substrate/product and cofactor (Fig. 4A and B). In AurF, the 4-nitrobenzoic acid (pNBA) product is held over Fe1 by polar contacts between the product carboxylate and side chains of Tyr93 and Asn200 from core helices α1 and α3, respectively. In the Fe_2_(II/II)-SznF•L-NMA model, the approach and number/type of contacts are similar. Our model predicts that the substrate is held over Fe1 by the aforementioned interactions with its α-ammonium and α-carboxylate groups. The FDO *N*-oxygenases have also resisted characterization in their substrate-bound forms, perhaps due to relatively low substrate affinities. In both the FDO and HDO systems, the failure of the substrate to impact the kinetics of O_2_ addition may indicate that this step precedes substrate binding in the preferred pathway (18, 21, 39). In this scenario, the Fe_2_(II/II)-SznF•L-NMA complex would be out of sequence and challenging to capture in the crystal.

### Bioinformatic analysis of the HDO superfamily

Comparison of the known structural and functional characteristics of SznF and UndA highlights potentially predictive correlations. The presence of the carboxylate ligand from aux α2 may (i) obviate direct substrate coordination to the cofactor and thus its triggering of O_2_ addition and (ii) stabilize the diiron-O_2_ adduct with respect to O-O-bond cleavage, favoring oxidation only of substrates that can attack as nucleophiles. Conversely, the absence of the extra carboxylate should leave this site vacant for substrate coordination, triggering of O_2_ addition, and (more speculatively) progression to a more reactive complex capable of H• abstraction from an aliphatic carbon (e.g., C3 of lauric acid, as proposed in recent studies of UndA). We searched sequence databases to segregate potential HDOs into groups that conserve the aux α2 β-turn insertion harboring the extra Glu from those that don’t. Use of an alignment score of 51 in generating a sequence similarity network (SSN) from the ∼ 9600 predicted HDO proteins that we identified (Fig. 5) separated the reference sequences [SznF and RohS (validated *N*-oxygenases), UndA and BesC (validated desaturases), and CADD (unknown function)] into different clusters, consistent with their distinct activities and targets. Within the top 80 clusters in the network (ranked by number of sequences), we searched for conservation of an aux α2 β-turn and coordinating carboxylate by manual alignment to SznF and other reference sequences. Predicted protein sequences in eight of these clusters (clusters 1, 7, 9, 11, 17, 22, 70, and 76) conserve features of the insertion/extra-ligand motif (Fig S17, Table S2). Clusters 9 and 11 contain the known *N*-oxygenases, RohS and SznF, respectively. We suspect that the other six clusters might also contain *N-*oxygenases with properties similar to SznF. One of these groups, cluster 1, is the largest of the SSN, containing 1780 sequences, roughly 20% of the HDO sequence pool evaluated here. Cluster 1 is enriched in sequences from non-pathogenic *Streptomyces* and *Mycobacterium* organisms (Fig. S17). Each of the remaining four clusters contains 50-200 sequences, and these are similarly dominated by predicted proteins from environmental aquatic and soil microbes, as opposed to pathogens or commensal bacteria. Apart from cluster 11, in which all sequences are sufficiently long (Fig. S18) to suggest fusion to a separate (e.g., cupin) domain, the remaining predicted *N-*oxygenases are uniformly shorter, indicating that they likely act as simple *N*-oxygenases. Testing of these top-level functional predictions is likely to vastly expand the known repertoire of the emerging HDO superfamily.

**Fig. 5.**
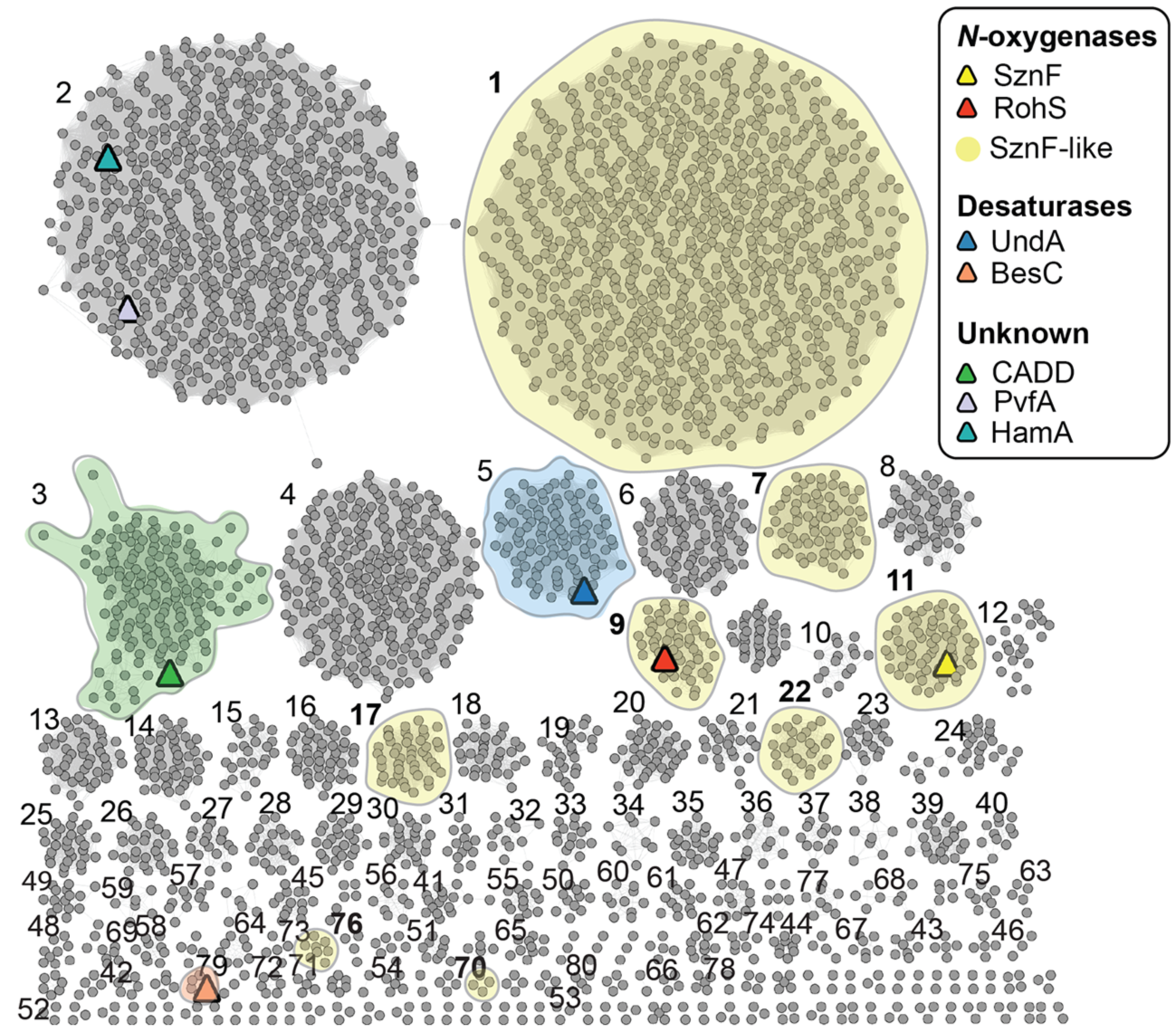
Bioinformatic analysis of the HDO superfamily. A sequence similarity network of 9600 predicted HDO sequences shows that many sequence families remain uncharacterized at the structural and/or functional level. Homologs of known *N*-oxygenases (red, yellow), oxidative desaturases (blue, peach), or structurally characterized proteins of unknown function (green, lavender, teal) are indicated as a colored triangle. Clusters designated SznF-like are shaded yellow, indicating conservation of the extra Glu ligand in aux α2.

## Methods

Detailed materials and methods are provided in the *SI Appendix*.

### Stopped-flow absorption (SF-Abs) spectroscopy

Stopped-flow absorption experiments were performed at 5 °C on an Applied Photophysics Ltd. (Leatherhead, UK) SX20 stopped-flow spectrophotometer housed in an MBraun anoxic chamber and equipped with either a photodiode-array (PDA) or photomultiplier tube (PMT) detector. Details of reaction conditions and data analysis are provided in the figure legends and *SI Appendix*.

### X-ray crystallographic characterization of apo and Fe_2_(II/II)-SznF

Apo N-terminally His_6_-tagged SznF was overexpressed and purified as detailed in the *SI Appendix*. SznF was crystallized under anoxic conditions in a stringently apo form in 16% (w/v) PEG 4000, 0.1 M MgCl_2_, and 0.1 M Tris pH 8.5. To obtain the Fe(II)-bound form of the enzyme, crystals were soaked for 5-10 min in precipitant solution supplemented with 10 mM ferrous ammonium sulfate. Data collection and refinement procedures are described in the *SI Appendix*.

### Bioinformatic analysis of HDO enzymes

An initial pool of 42,581 unique sequences annotated as either heme-oxygenase or heme-oxygenase-like proteins was extracted from Interpro (Nov. 2019 release). Using length filters and a custom Python script, this pool was analyzed for conservation of the HDO core helix diiron ligands (Fig. S1) to obtain a final set of ∼ 9600 candidate HDO enzymes as input for SSN generation with the Enzyme Function Initiative Enzyme Similarity Tool (57). Using manual and automated sequence alignments and homology model generation, eight of the top 80 (ranked by number of sequences) sequence clusters were annotated as likely *N*-oxygenases on the basis of conservation of the additional carboxylate ligand and β-turn motif in aux α2.

## Acknowledgements

We gratefully acknowledge use of the resources of the Advanced Photon Source, a U.S. Department of Energy (DOE) Office of Science User Facility operated for the DOE Office of Science by Argonne National Laboratory under Contract No. DE-AC02-06CH11357. Use of the LS-CAT Sector 21 was supported by the Michigan Economic Development Corporation and the Michigan Technology Tri-Corridor (Grant 085P1000817). GM/CA@APS has been funded in whole or in part with Federal funds from the National Cancer Institute (ACB-12002) and the National Institute of General Medical Sciences (AGM-12006). The Eiger 16M detector at GM/CA-XSD was funded by NIH grant S10 OD012289. This work was additionally supported by National Institutes of Health grants GM119707 to A.K.B. and GM138580 to J.M.B.

