## Supporting Information for "Structure and assembly of the diiron cofactor in the heme-oxygenase-like domain of the *N*-nitrosourea-producing enzyme SznF"

*SI Appendix*

**SF-Abs experimental methods.** In all experiments, the Applied Photophysics SX20 stopped-flow apparatus was configured for single mixing. The path length was 1 cm, except in reactions of the SznF•Fe(II) complex experiments ferrozine, in which case it was 0.2 cm. Time-dependent absorption spectra (1000 data points collected over 100 s) were acquired with a logarithmic time base. At least three trials were performed for each reaction condition to verify reproducibility. The extremely high signal-to-noise ratio and reproducibility of individual traces resulting from the high absorptivity of the μ-peroxo-$Fe_2$(III/III) and Fe(II)•(ferrozine)$_3$ complexes rendered averaging of kinetic traces unnecessary.

*Reaction of the Fe(II)•SznF(-Δcupin) complex with $O_2$.* Reactant solutions containing Fe(II) and protein were prepared in the MBraun anoxic chamber. These solutions contained 0.3 mM SznF (wild-type or Δcupin variant) and 0-1.8 mM (0-6 molar equiv) Fe(II) in 20 mM sodium HEPES (pH 7.5), 50 mM NaCl, and 5% glycerol (*v/v*). The anoxic protein solution was mixed at 5 °C with an equal volume of the same buffer at equilibrium (at 5 °C) with ~ 1.05 atm $O_2$ (~1.8 mM). Absorbance was monitored with the photodiode array detector.

*Reaction of the Fe(II)•SznF(-Δcupin) complex with ferrozine.* The reactant Fe(II)•SznF(-Δcupin) complex was prepared similarly (in the anoxic chamber) and had 80 μM SznF and 80-400 μM Fe(II) in 50 mM sodium HEPES (pH 7.5), 50 mM NaCl, 5% glycerol (*v/v*) and 1 mM sodium dithionite. This solution was mixed with an equal volume of an anoxic solution containing 4 mM ferrozine in the same buffer. Absorbance at 562 nm ($A_{562}$) was monitored with the photomultiplier tube.

*Reaction of apo SznF(-Δcupin) simultaneously with Fe(II) and $O_2$.* In these experiments, an $O_2$-saturated solution of 0.30 mM apo SznF (wild-type or Δcupin variant) in 50 mM sodium HEPES (pH 7.5), 50 mM NaCl, and 5% glycerol (*v/v*) was mixed with an equal volume of an anoxic solution of 0-1.8 mM Fe(II) in the same buffer. Absorbance was monitored with the photodiode array.

*Reaction of the $Fe(II)_{0.5\text{-}1}$•SznF(-Δcupin) complex with additional Fe(II) and $O_2$.* In these experiments, the anoxic protein reactant solution contained 0.30 mM SznF (wild-type or Δcupin variant) and either 0.30 mM (for wild-type SznF) or 0.15 mM (for the Δcupin variant) Fe(II) in 100 mM sodium HEPES (pH 7.5), 50 mM NaCl, and 5% glycerol (*v/v*). This solution was mixed at 5 °C with an equal volume of an $O_2$-saturated solution containing either 1.5 mM Fe(II) (for wild-type SznF) or 0.9 mM Fe(II) (for the Δcupin variant) in 1 mM $H_2SO_4$ (to stabilize the reduced iron toward $O_2$). Absorbance was monitored with the photodiode array.

**SF-Abs Data Analysis.** *Physical model and equations used in regression analysis of the SF-Abs experiments monitoring the μ-peroxo-$Fe_2$(III/III) intermediate.* The physical model invoked to rationalize the kinetics of formation and decay of the *μ*-peroxo-$Fe_2$(III/III) intermediate in SznF is illustrated in Scheme S1. The protein is a dimer, and the HDO domain of each monomer has a dinuclear site. An Fe(II):SznF ratio of one (two per dimer) appears sufficient to completely convert the apo protein (black state at left) to the complex with one of two subsites of the diiron site (the tighter or more avid one) filled in each monomer (all blue complex). Increasing Fe(II):SznF ratios allow for increasing association in the less avid subsite of one of the two monomers (mixed blue and red complex). Filling of the fourth site (all red complex) appears to be disfavored, even at the maximum iron equivalency tested [Fe(II):SznF = 6; Fe(II) = 1.8 mM]. This low effective affinity is likely a result of an anti-cooperative interaction across the dimer, as has previously been seen in the homodimeric ferritin-like diiron β subunit of *Escherichia coli* class Ia ribonucleotide reductase (1, 2). In the anoxic preincubation of SznF(-Δcupin) with ≥ 1 equiv Fe(II), a distribution of mononuclear and dinuclear complexes (all red, all blue, and red/blue complexes) are present. Mixing with $O_2$ initiates conversion to the 629-nm-absorbing *μ*-peroxo-$Fe_2$(III/III) intermediate (PXO). Monomers with both subsites filled (Scheme S1B, DFS, red) can react with $O_2$ directly, thus supporting a two-step formation-and-decay sequence, whereas monomers with only the more avid subsite filled (MFS, blue) must first bind Fe(II) in the less avid subsite and then react with $O_2$, thus leading to a three-step sequence. The $A_{629}$ kinetic traces were fit by Eq. 1, which is appropriate for this physical model and was adapted from a previously published derivation (3). In this equation, the ε terms are the molar absorption coefficients at 629 nm of the four species in the mechanism, and the $k$ terms are rate constants, as defined in Scheme S1B. The Fe(II)-containing complexes are effectively transparent ($\varepsilon_{629}$ ~ 0), whereas Iterative analysis indicated that the diferric product (DFC) contributes 0.039 times the absorbance of the $\mu$-peroxo-$Fe_2$(III/III) complex at 629 nm. This dependency was explicitly incorporated in the fitting analysis to reduce parameter space. The blue and red terms in Eq. 1 account for the kinetics of the reactions of the MFS (blue) and DFS (red) complexes, respectively. Their initial concentrations ($[MFS]_0$ and $[DFS]_0$), which dictate the amplitudes of the fast and slow rise phases of $A_{629}$ in Figure 1A-B, were treated as adjustable parameters in the fit, and the relative proportions of each complex at each Fe(II):SznF(-Δcupin) are plotted in Fig. S6A,C. The corresponding rate constants from the fits are plotted in Fig. S6B,D.

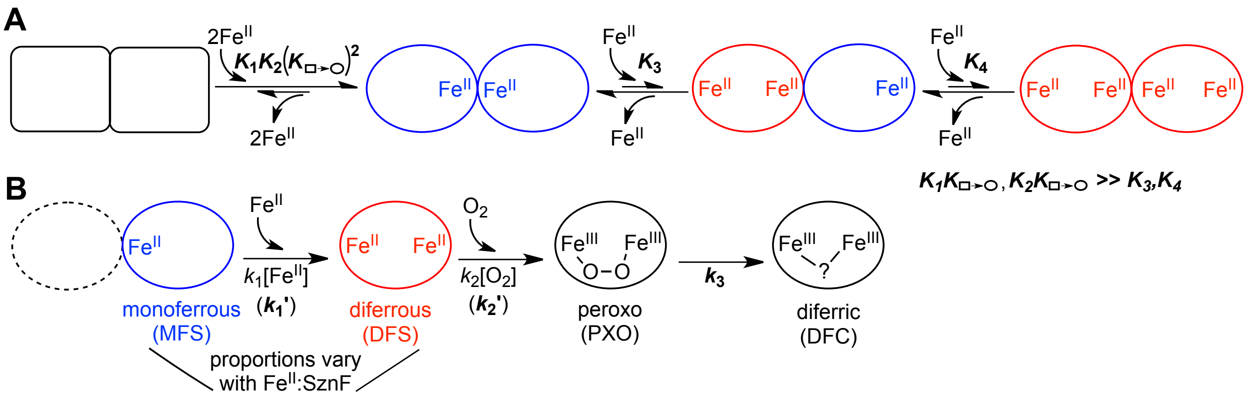


**Scheme S1.** Physical model considered in analysis of the kinetics of formation and decay of the $\mu$-peroxo-$Fe_2$(III/III) intermediate in SznF.

$$A_\lambda(t) = [\mathrm{MFS}]_0\left\{\varepsilon_{\mathrm{DFC}} + (\varepsilon_{\mathrm{MFS}} - \varepsilon_{\mathrm{DFC}})e^{-k_1' t} + (\varepsilon_{\mathrm{DFS}} - \varepsilon_{\mathrm{DFC}})\left(\frac{k_1'}{k_2' - k_1'}\right)\left(e^{-k_1' t} - e^{-k_2' t}\right) + (\varepsilon_{\mathrm{PXO}} - \varepsilon_{\mathrm{DFC}})\left(\frac{k_2' k_1'}{k_2' - k_1'}\right)\left\{\left(\frac{1}{k_3 - k_1'}\right)\left(e^{-k_1' t} - e^{-k_3 t}\right) - \left(\frac{1}{k_3 - k_2'}\right)\left(e^{-k_2' t} - e^{-k_3 t}\right)\right\}\right\} + [\mathrm{DFS}]_0\left\{\varepsilon_{\mathrm{DFC}} + (\varepsilon_{\mathrm{DFS}} - \varepsilon_{\mathrm{DFC}})e^{-k_2' t} + (\varepsilon_{\mathrm{PXO}} - \varepsilon_{\mathrm{DFC}})\left(\frac{k_2'}{k_3 - k_2'}\right)\left(e^{-k_2' t} - e^{-k_3 t}\right)\right\} \quad \textbf{Eq. 1}$$

In the experiments in which apo SznF was reacted simultaneously with Fe(II) and $O_2$ and those in which a solution of partially iron-loaded protein was mixed with Fe(II) and $O_2$, the equation used in the fitting analysis was essentially just the blue terms in Eq. 1. For the case of the experiments with apo SznF, use of this equation approximates the several steps needed to form the $\mu$-peroxo-$Fe_2$(III/III) complex (binding of iron in the more avid site, the protein conformational change, binding of iron in the less avid site, and trapping of $O_2$) as just two steps. These four steps have not been explicitly kinetically resolved in the experiments in this study (but it may be possible to do so in the future by sequential-mixing experiments incorporating varying orders of addition and reaction times between mixes), and so approximation of the multiple steps as a two-step sequence avoided unwarranted degrees of freedom (i.e., too many parameters unconstrained by relevant data) in the regression analysis. Across all fits, the optimized rate constant for decay of the $\mu$-peroxo-$Fe_2$(III/III) intermediate was 0.3 ± 0.15 $s^{-1}$, in agreement with the value of 0.34 $s^{-1}$ reported in the previous study.

*Regression analysis of the $A_{562}$ kinetic traces from reactions of the Fe(II)•SznF(-Δcupin) complex with ferrozine.* The objective of this analysis was to measure the amplitudes of the fast and slow phases of chelation in order to estimate the concentrations of free and bound Fe(II) as a function of Fe(II):SznF(-Δcupin) ratio. This aim was achieved by fitting the traces by the equation for multiple (2 or 3) parallel, first-order processes. The slower phases of color development involve a minimum of two sequential steps: dissociation from SznF and chelation by ferrozine. In addition, dissociation of Fe(II) from multivalent complexes could also involve a sequence of two or more steps [e.g., loss of Fe(II) from the less avid subsite of the dinuclear site in the HDO domain followed by dissociation from the tighter subsite]. However, as the point of the analysis was a qualitative understanding of the concentrations of free and bound iron, the use of multiple exponentials, corresponding to multiple parallel processes, was deemed adequate.

**X-ray crystal structure solution of apo- and $Fe_2$(II/II)-SznF.** N-terminally $His_6$-tagged apo SznF was overexpressed and purified by affinity chromatography as previously described (4), but with an additional purification step that was necessary for protein crystallization. Prior to crystallization trials, protein samples were chromatographed at 4 °C on a HiLoad 16/600 Superdex 200 gel filtration column controlled by an ÄKTA Pure FPLC system (GE Healthcare). The mobile phase was 20 mM sodium HEPES (pH 7.5) buffer at 0.5 mL/min. Protein samples were flash-frozen in liquid $N_2$ for long-term storage at -80˚C. Prior to crystal tray setup, the protein was thawed and kept open in a COY anoxic chamber for at least 2 h. SznF was crystallized in its apo form by using the hanging drop vapor diffusion method. Protein samples were mixed with an equal volume of a precipitant solution containing 16% (w/v) PEG 4000, 0.1 M $MgCl_2$, and 0.1 M Tris-HCl, pH 8.5. Crystals appeared within three days and grew to full size within a week. Prior to harvest, crystals were soaked for 5-10 min in precipitant solution supplemented with ferrous ammonium sulfate [10 mM final Fe(II)] diluted in the well solution. The Fe(II) soak solution was prepared from an acidic stock [0.20 M Fe(II) in 0.1 M $H_2SO_4$] and diluted to a final acid concentration of 5 mM $H_2SO_4$. Crystals were harvested in rayon loops and flash frozen in liquid $N_2$ without any additional cryoprotectant. Crystals of the apo protein were prepared identically but with the Fe(II) soak step omitted and all steps performed aerobically.

Crystallographic datasets were collected at the Life Sciences Collaborative Access Team (LS-CAT) and the National Institute of General Medical Science and National Cancer Institute Collaborative Access Team (GM/CA-CAT) beamlines at the Advanced Photon Source (Argonne National Laboratory, Argonne, IL). All diffraction datasets were processed with the HKL2000 package (5). Phases were obtained by molecular replacement (MR) with PHASER (6) implemented within the CCP4 software package (7). A previously solved structure of SznF (PDB accession code 6M9R) was used as the MR input model. Model building was carried out in Coot (8). Refinement procedures were performed in Refmac5 (9) and Phenix (10). Coordinates were analyzed for Ramachandran outliers and other geometric parameters with the Molprobity server

(11). Cavity analysis was performed with Hollow (12) and conservation analysis was accomplished by using the ConSurf server (13-17). Figures were prepared with the PyMOL molecular graphics software package (Schrödinger, LLC).

Crystals of both apo-SznF and $Fe_2$(II/II)-SznF belong to the $P2_1 2_1 2_1$ space group with two monomers in the asymmetric unit. The final model of apo-SznF consists of residues 4-132, 138-150, 157-471 in chain A; 4-133, 137-150, 157-471 in chain B; and 592 water molecules. In the model of $Fe_2$(II/II)-SznF, chain A consists of residues 1-132, 136-150, and 157-471 with two additional residues at the N-terminus from the $His_6$-affinity tag; chain B consists of residues 4-133, 136-150, and 157- 471. The final model also contains six Fe(II) ions and 411 water molecules. In each chain, one Fe(II) was modeled in the C-terminal cupin domain and the remaining two Fe(II) ions were found in the central HO-like domain. In the C-terminal cupin domain, Fe(II) occupancies were 100% and 97%, respectively, for chains A and B. The Fe 1 site occupancies were modeled as 93% and 100%, respectively, for chains A and B. The Fe 2 site occupancies were modeled as 67% and 84%, respectively, for chain A and B.

**Bioinformatic analysis of HDO enzymes.** A pool of 42,581 unique sequences annotated as heme-oxygenase or heme-oxygenase-like proteins in three different superfamilies (IPR016084, IPR039068, IPR002051) was obtained from Interpro 77.0 (November 2019 release) (18). A length filter was applied to remove all sequences less than 100 and greater than 600 residues. In a process automated by a custom Python script, the remaining 40,590 candidates were aligned in pairwise fashion with the sequences of five reference proteins – SznF, UndA, CADD, RohS, and BesC (4, 19-26) (Table S3) – to test for conservation of the iron ligands of the reference proteins in the identified sequences. Sequences were examined for only the five ligands in the H-$X_n$-E-$X_n$-H-$X_3$-D/E/N-$X_2$-H (n = variable gap) pattern spanning core helices 1 and 3 of the reference proteins (Fig. S1); the isolated carboxylate ligand in core helix 2 proved challenging to identify unequivocally, because of low similarity and insufficient alignment constraints. To evaluate conservation of the ligand pattern in a given protein identified in the initial similarity search, we compared not only the position nominally aligned with the known ligand but also the two flanking amino acids in both directions. If one of these five residues was a potential ligand (D/E/H/N), we considered it as conservative of this element of the query pattern. If none of these five consecutive positions in a target protein contained a potential ligand, we considered it to violate the ligand pattern and discarded the sequence. This analysis yielded a final pool of ~9600 unique predicted HDO sequences.

A FASTA format file containing these sequences was used as input for generation of a sequence similarity network (SSN) with the Enzyme Function Initiative Enzyme Similarity Tool (EFI-EST) (27-30). An alignment score of 51, corresponding to 40% sequence identity, was used as a threshold for separating sequences into distinct clusters. The resulting file was analyzed in Cytoscape (31). The top 40 sequence clusters were analyzed for sequence length, number of sequences per node, and conservation of the SznF Glu189 equivalent. The latter property was investigated both manually by multiple sequence alignment in ClustalX (32) to one or more selected reference sequences and in an automated fashion with a second custom Python script. In sequence clusters that appeared to conserve the seventh carboxylate ligand, we additionally generated homology models (33-35) to further validate this prediction.

**Molecular dynamics (MD) simulation of the L-NMA•$Fe_2$(II/II)-SznF complex**. A model of the substrate, L-NMA, was generated in JLigand (36), implemented from within the CCP4 software package (7). Model constraints (.lib file) for L-NMA were generated with the PRODRG server (37) by using the output from JLigand as the input file for PRODRG. Output files from PRODRG (.pdb and .lib files) were used for manual substrate docking in Coot, guided by our hypotheses about binding interactions gleaned from the SznF structure, and merged with the $Fe_2$(II/II)-SznF protein coordinates (8). The L-NMA•SznF complex was prepared for molecular dynamics (MD) simulations using the Xleap program in AmberTools (38) with the ff14SB forcefield (39). The complex was solvated in TIP3P water (40) using a 10 Å octahedral box. $Na^+$ and $Cl^-$ ions were added first to neutralize and then to achieve a final concentration of 150 mM NaCl. Parameters for L-NMA were obtained using Antechamber (41, 42) in AmberTools. All minimizations and production runs were performed with Amber18 (38, 43). Minimization was achieved in multiple rounds via the following protocol: 5000 steps of steepest descent and 5000 steps of conjugate gradient, with i) 500 kcal/mol-Å$^2$ restraints on all solute atoms, ii) 100 kcal/mol- Å $^2$ restraints on all solute atoms, iii) 100 kcal/mol-Å$^2$ restraints on the substrate and Fe(II) ions, iv) all restraints removed. Following minimization, the complex was heated from 0 to 300 K using a 100-ps run, constant volume periodic boundaries and 10-kcal/mol•Å$^2$ restraints on all solute atoms. Equilibration was performed in three phases: i) 10 ns MD with 10-kcal/mol•Å$^2$ restraints on all solute atoms, ii) 10 ns with 1-kcal/mol•Å$^2$ on all solute atoms, iii) 10 ns with 1-kcal/mol•Å$^2$ restraints on Fe(II) and substrate atoms. Finally, an unrestrained MD simulation was performed, allowing a conformational sampling of the substrate in the active site.

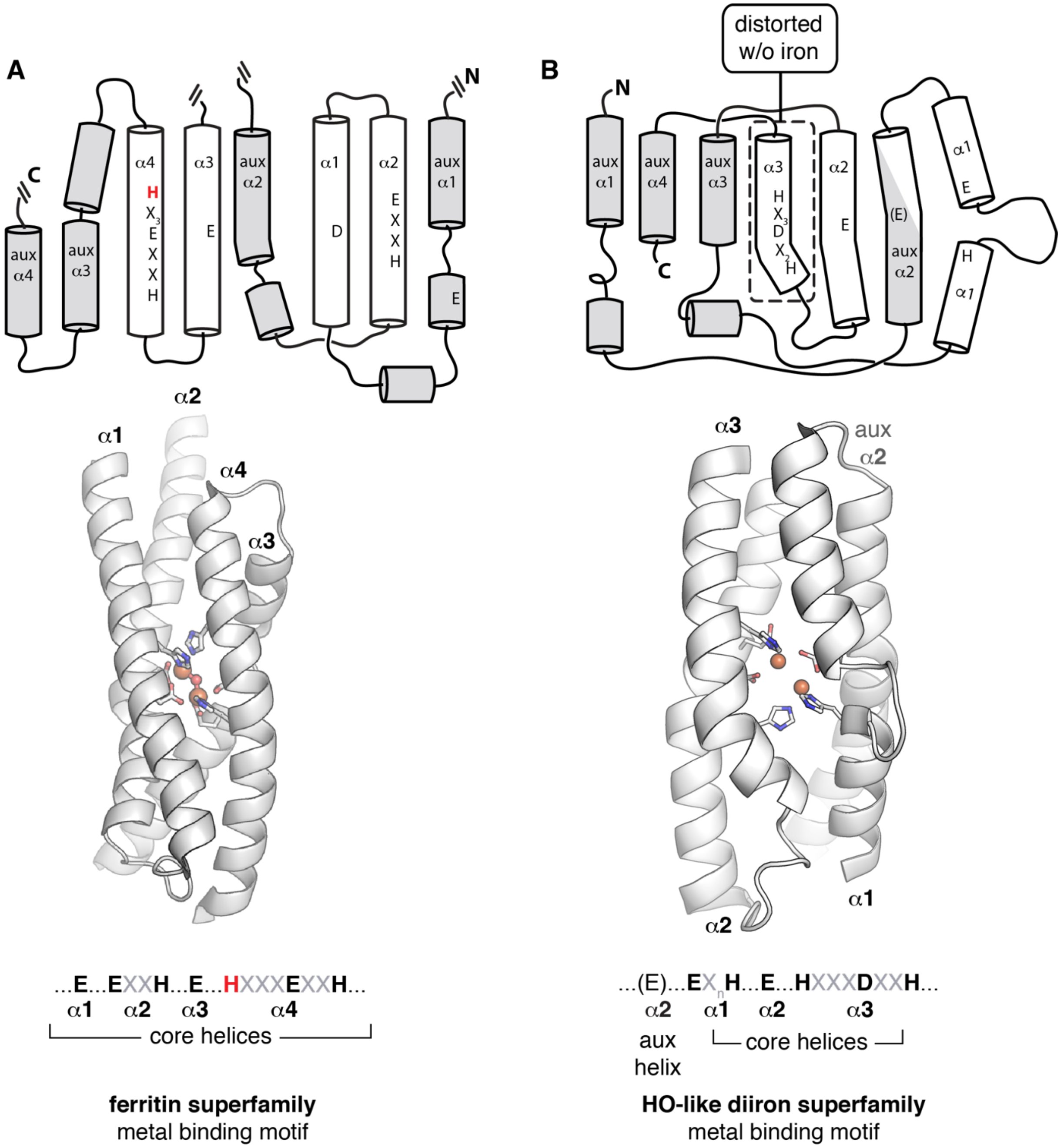


**Fig. S1.** Comparison of the topology and core metal binding helices in ferritin superfamily dimetal oxygenases/oxidases (FDOs) and heme-oxygenase superfamily diiron oxygenases/oxidases (HDOs). (*A*) FDOs use two pairs of regular α-helices to provide His and carboxylate ligands to their dimetal cofactors. In the FDO *N*-oxygenases, a seventh ligand, an additional His (red text, bottom panel), is contributed by the C-terminal core helix, α4. (*B*) HDOs typically use three distorted or interrupted helices to provide a different pattern of His and carboxylate ligands. The structure of $Fe_2^{II/II}$-SznF, an HDO *N*-oxygenase, reveals a seventh ligand contributed by an auxiliary helix, aux α2.

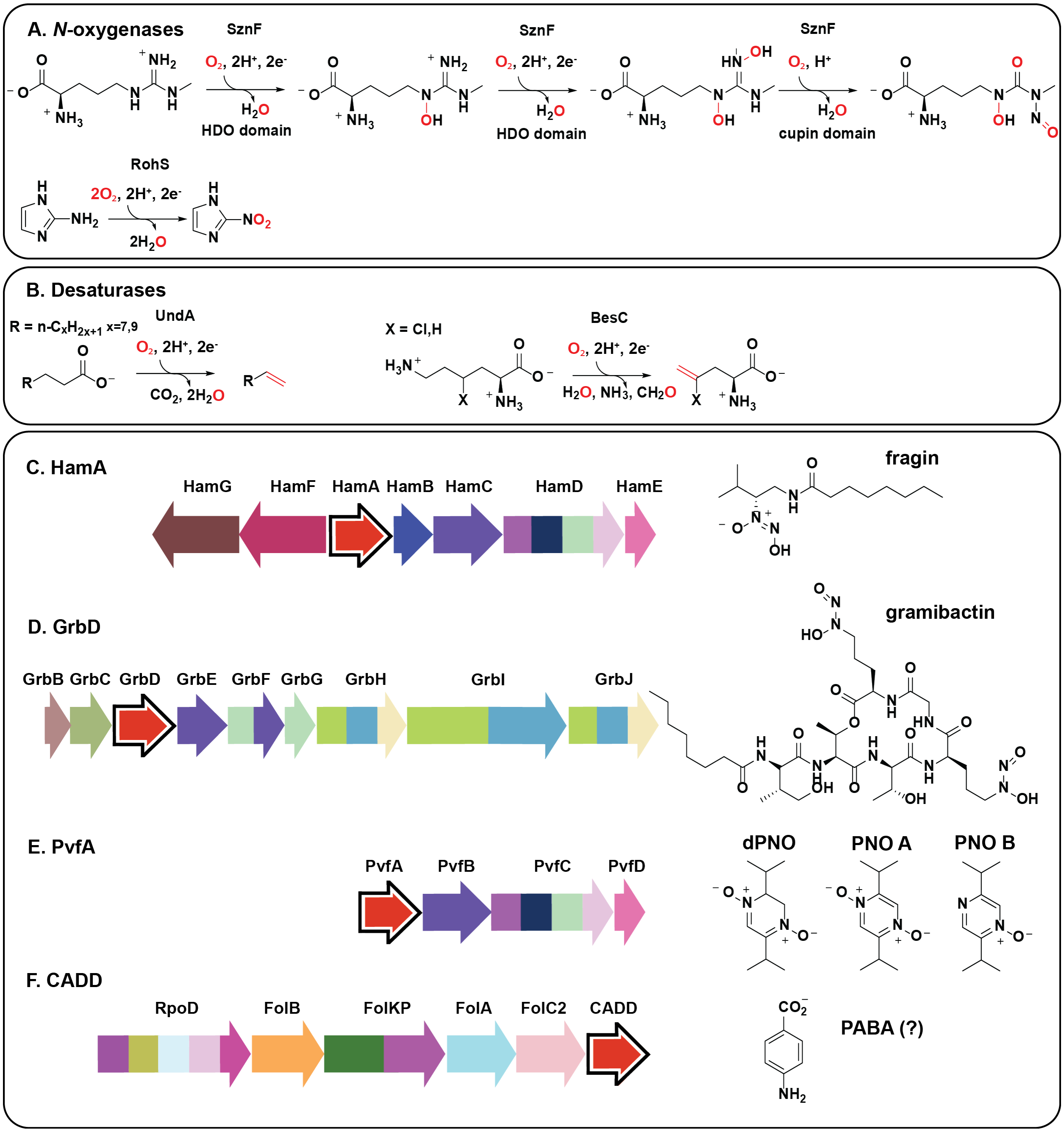


**Fig. S2.** Known and posited functions of HDOs. The known HDOs have either (*A*) *N*-oxygenase or (*B*) desaturase-lyase activities. Biosynthetic gene clusters encoding proposed HDOs (*C*) HamA, (*D*) GrbD, (*E*) PvfA, and (*F*) CADD. Putative HDOs are shown in red, and known or proposed products of the biosynthetic clusters are shown at right. HamA is a heme-oxygenase-like protein encoded adjacent to cupin protein HamB (44). GrbD is implicated in gramibactin biosynthesis and has sequence similarity to SznF (45, 46). The *pvf* gene cluster is homologous to the *ham* gene cluster (47), and, interestingly, each encodes an additional FDO *N*-oxygenase (HamC, PvfB), suggesting that the HDO component might have an alternative function. The activity of CADD is not yet known (24, 48).

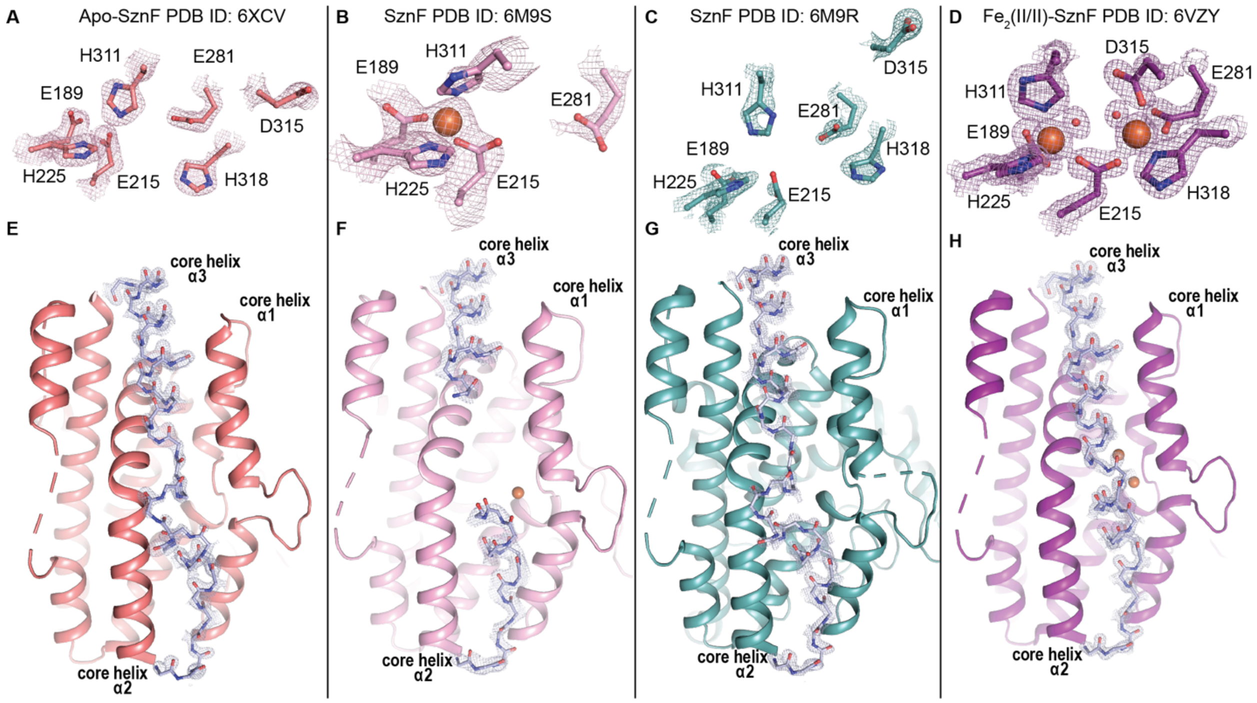


**Fig. S3.** Comparison of the HDO active site and overall structure across all SznF structures solved to date. The apo SznF structure solved in this study (*A*) (*salmon*, 6XCV) shows interpretable electron density for all metal-binding side chains, but some are oriented away from the active site. Previously published structures of SznF that had been exposed to Fe(II) and $O_2$ (*B*) (*pink,* 6M9S) and (*C*) (*light teal*, 6M9R) showed weaker electron density for core $\alpha$3, consistent with heterogeneity and/or disorder. The structure of $Fe_2^{II/II}$-SznF solved in this study (*D*) (*purple,* 6VZY) shows both Fe sites highly occupied and all ligands oriented toward the cofactor. Ligands are shown in stick format. $2F_o$-$F_c$ electron density maps contoured to either 1.0$\sigma$ (6XCV, 6M9S, 6M9R) or 1.5$\sigma$ (6VZY) are shown in colored mesh. Iron ions and water molecules are shown as orange and red spheres, respectively. (*E-H*) Cartoon representation of the HDO domain with core $\alpha$3 backbone shown in stick format (highlighted in *light blue* with $2F_o$-$F_c$ electron density maps contoured to 1.5$\sigma$). (*E*) Apo SznF (*salmon*, 6XCV) shows backbone density for all residues in core $\alpha$ helix 3 despite helical disorder. (*F*) In several of the structures of Fe(II)/$O_2$-treated SznF solved previously (*light pink*, 6M9S), a gap of 9 residues remained unmodeled in core $\alpha$3. (*G*) In structures from other datasets (*light teal*, 6M9R), more nearly contiguous electron density for the backbone in core $\alpha$ helix 3 was seen, but density for the side chains was weak, suggesting disorder. (*H*) The structure of $Fe_2$(II/II)-SznF (*purple*, 6VZY) solved in this study shows full ordering of the core $\alpha$3 helix.

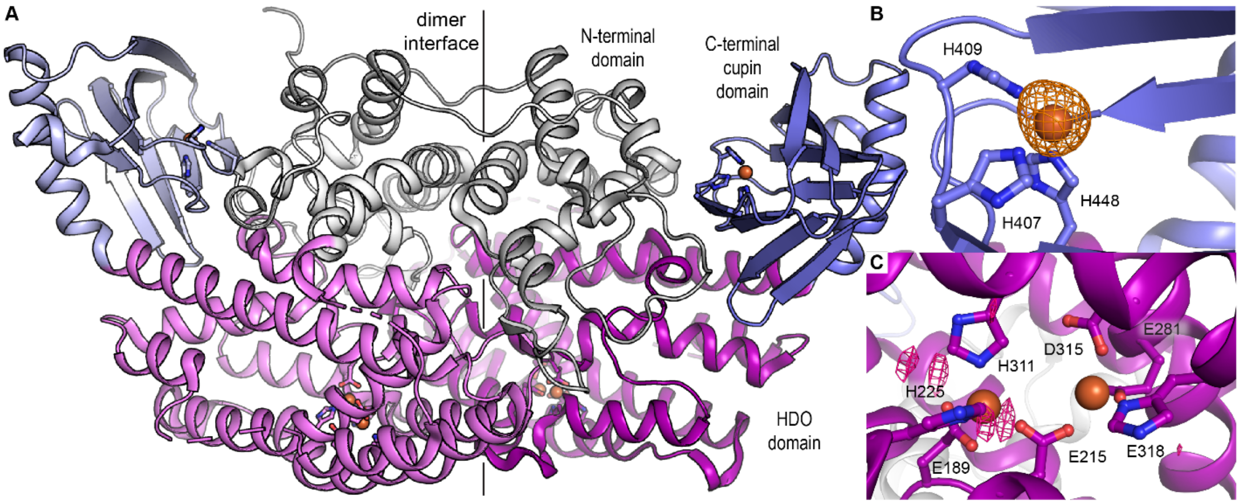


**Fig. S4.** Cartoon representation of the entire SznF homodimer and zoomed in view of the cupin and HDO active sites. (*A*) A ribbon diagram of the homodimer from the structure of the protein with the HDO and cupin domains loaded with Fe(II) (6VZY). The N-terminal domain is shown in *grey/white*, the central 7-helical-bundle HDO domain is shown in *purple/light purple*, and the C-terminal cupin domain is shown in *blue/light blue*. (*B*) A zoomed-in view of the cupin domain cofactor site. An iron anomalous difference map is shown in *orange mesh* and contoured to 3.0$\sigma$. (*C*) A zoomed-in view of the active site in the HDO domain (*purple/light purple*) with metal binding ligands in *stick format* and Fe atoms shown as *orange spheres*. Apo SznF samples were prepared in M9 medium with Mn(II) supplementation and metal chelation after protein purification. To determine whether residual Mn(II) remained in the HDO metal binding site after this process, an Mn anomalous difference map was obtained, shown in hot pink mesh contoured to 3$\sigma$. Very little density is observed above the noise level, suggesting little to no occupancy of the HDO domain by Mn(II). Selected side chains are shown in *stick format* and iron ions are shown as *orange spheres*.

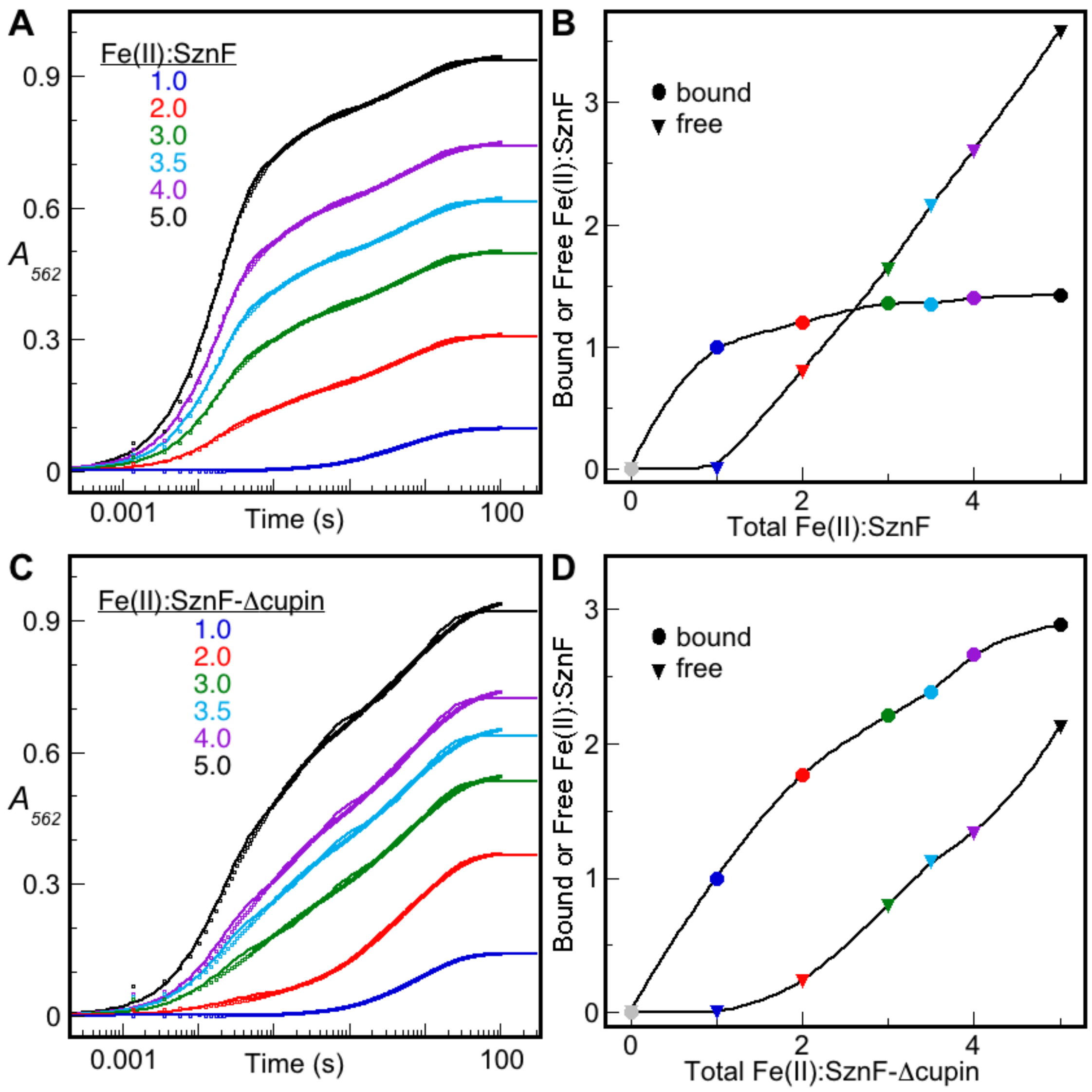


**Fig. S5.** Binding and dissociation of Fe(II) monitored by the kinetics of its chelation by the colorimetric indicator, ferrozine. An anoxic solution of 80 μM SznF (*A*, *B*) or SznF-Δcupin (*C*, *D*) containing the indicated molar ratio of Fe(II) was mixed with an equal volume of an anoxic solution of 4 mM ferrozine, and absorbance at 562 nm ($A_{562}$) was monitored as a function of reaction time. The traces are representative of at least 3 trials (SF "shots") for each condition. To estimate the concentration of free (rapidly chelating) and bound (slowly chelating) Fe(II), each trace was fit by the equation for two [Fe(II):SznF(-Δcupin) = 1] or three [for all other Fe(II):SznF(-Δcupin) ratios] parallel, first-order growths. The fractional amplitude of the fast phase and the summed fractional amplitudes of the two slower phases were multiplied by the initial Fe(II):SznF(-Δcupin) ratio for each experiment (trace), and the resultant equivalencies of free (*triangles*) and bound (*circles*) Fe(II) were plotted versus Fe(II):SznF(-Δcupin) ratio (*C* and *D*).

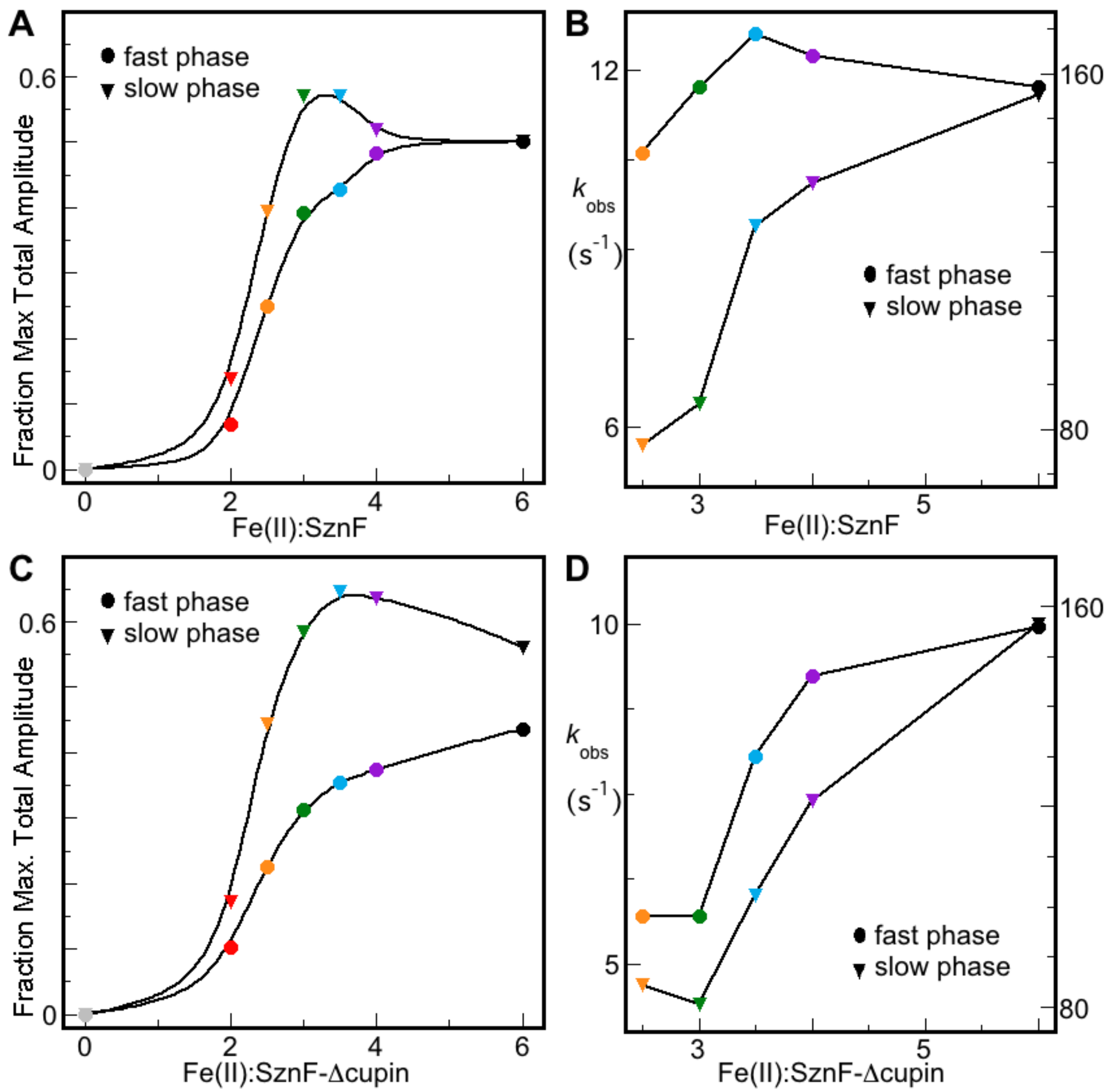


**Fig. S6.** Amplitudes and rate constants derived from regression analysis of the traces in main text Figure 1A-B according to Eq. 1 appropriate for Scheme S1. Panels A and B depict the results from analysis of the traces in Figure 1A, and panels C and D depict the results from analysis of the traces in Figure 1B.

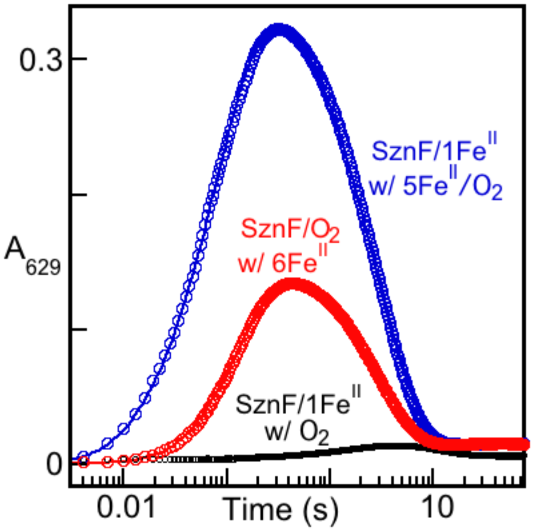


**Fig. S7.** $A_{629}$-versus-time traces monitoring the kinetics of the $\mu$-peroxo-$Fe_2$(III/III) intermediate after mixing of an anoxic solution of SznF containing 1 molar equiv Fe(II) with an equal volume of a solution containing either only $O_2$ (*black*) or $O_2$ with 5 additional equiv Fe(II) (*blue*). The trace obtained after direct mixing of apo SznF with all 6 equiv Fe(II) is shown for comparison (*red*).

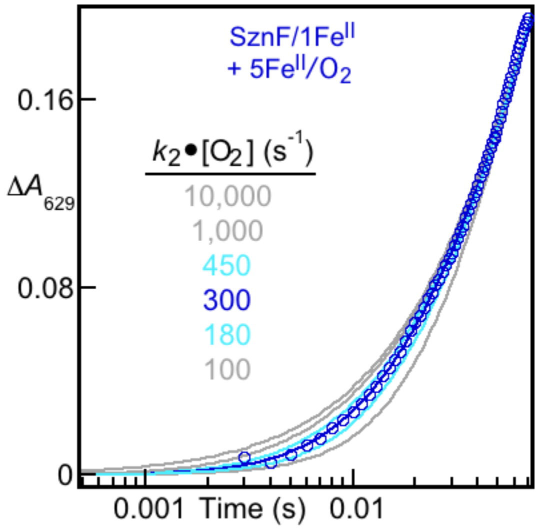


**Fig. S8.** Zoomed-in view of the early part of the $A_{629}$ kinetic trace obtained after mixing SznF containing 1 equiv bound Fe(II) simultaneously with 5 additional equiv Fe(II) and $O_2$ (Fig. S7, *blue*) to illustrate estimation of the rate constants for the Fe(II)- and $O_2$-addition steps comprising formation of the $\mu$-peroxo-$Fe_2$(III/III) complex. The solid lines are best-fit traces with varying effective first-order rate constants for the $O_2$-addition step. The effective first-order state constant for the Fe(II)-addition step – treated as an adjustable parameter in the fitting – was 11.8 ± 1.0 $s^{-1}$.

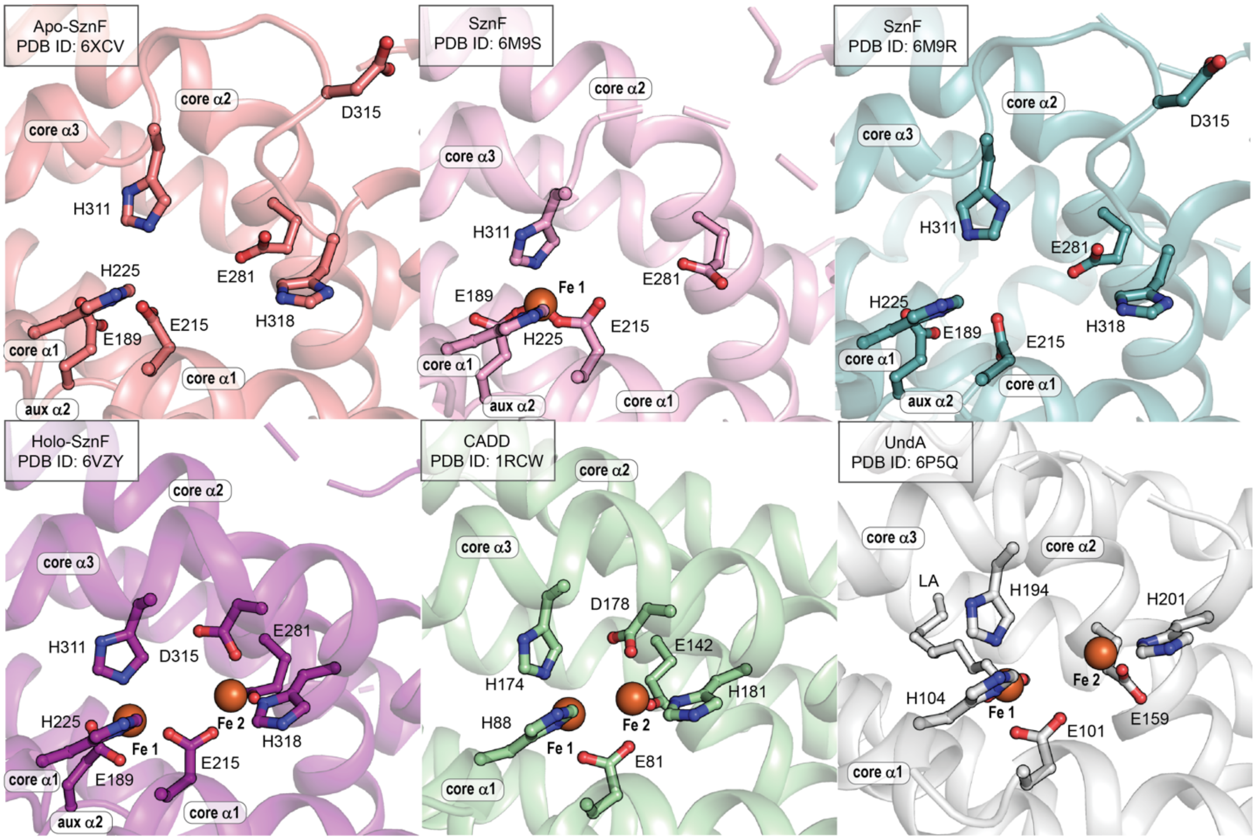


**Fig. S9.** Zoomed-in views of the metal-binding site in all HDOs that have been structurally characterized by x-ray crystallography. Structures of Apo and reconstituted SznF are shown in the top panels. Structures of $Fe_2$(II/II)-SznF (*left*), CADD (*center*) and UndA (*right*) are depicted in the bottom panels. In the structure of UndA, density for lauric acid (LA) substrate and two iron ions was seen, but site 2 had low occupancy and core α3 remained disordered.

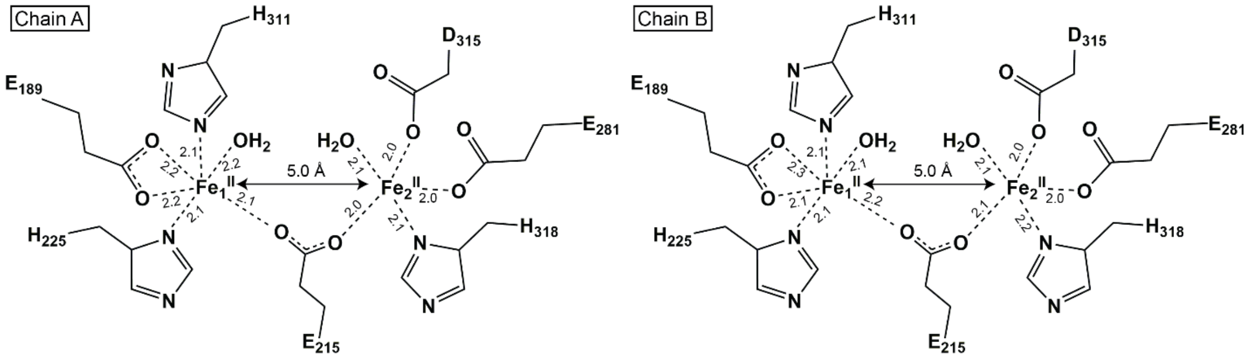


**Fig. S10**. Schematic diagram of the first coordination spheres in the active sites of the HDO domains in both monomers of the $Fe_2$(II/II)-SznF homodimer (chains A and B). Fe1 is coordinated by E189, E215, H225, and H311 and Fe site 2 is ligated by E215, E281, D315, and H315. Metal-metal and metal-ligand distances are shown in units of angstroms. The distances are very similar in the two monomers.

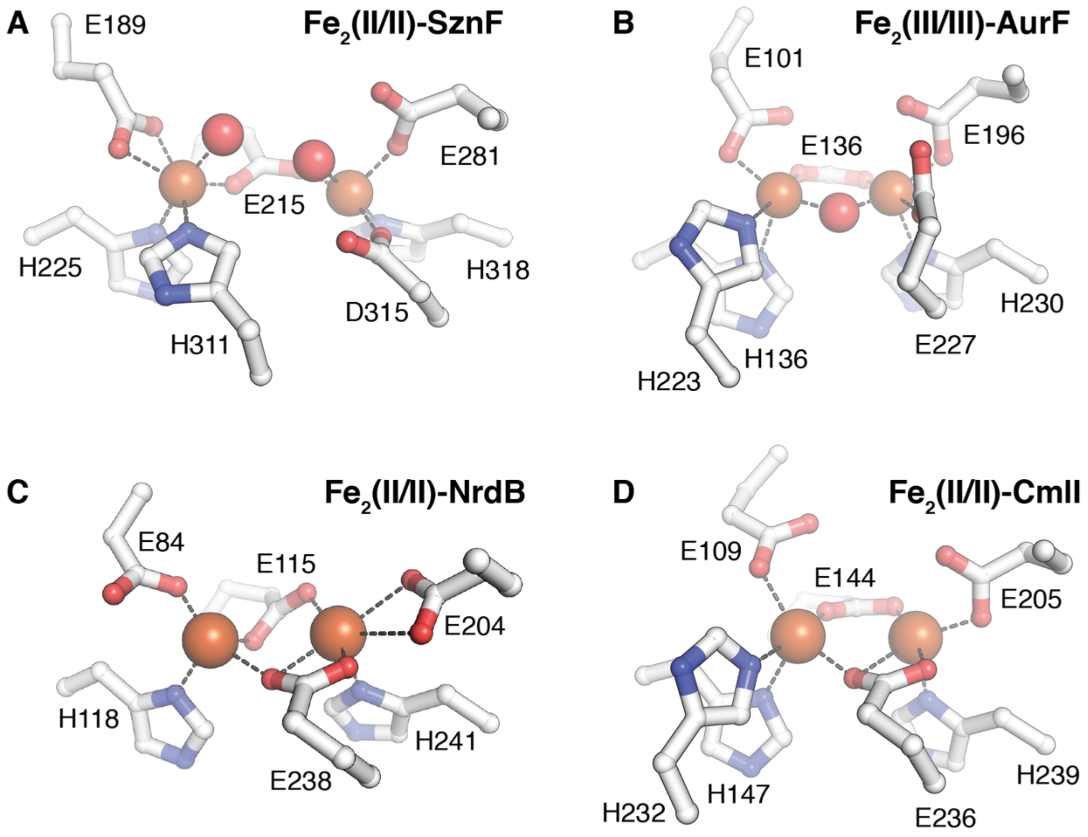


**Fig. S11.** Comparison of the metal-binding sites in the new structure of SznF with its HDO cofactor bound to those seen in representative structures of oxidized and reduced FDOs. (*A*) HDO *N*-oxygenase $Fe_2$(II/II)-SznF (PDB accession code 6VZY), (*B*) FDO *N*-oxygenase $Fe_2$(III/III)-AurF (PDB accession code 3CHI), (*C*) FDO class Ia ribonucleotide reductase β subunit $Fe_2$(II/II)-NrdB (PDB accession code 1PIY), (*D*) FDO *N*-oxygenase $Fe_2$(II/II)-CmlI (PDB accession code 5HYH).

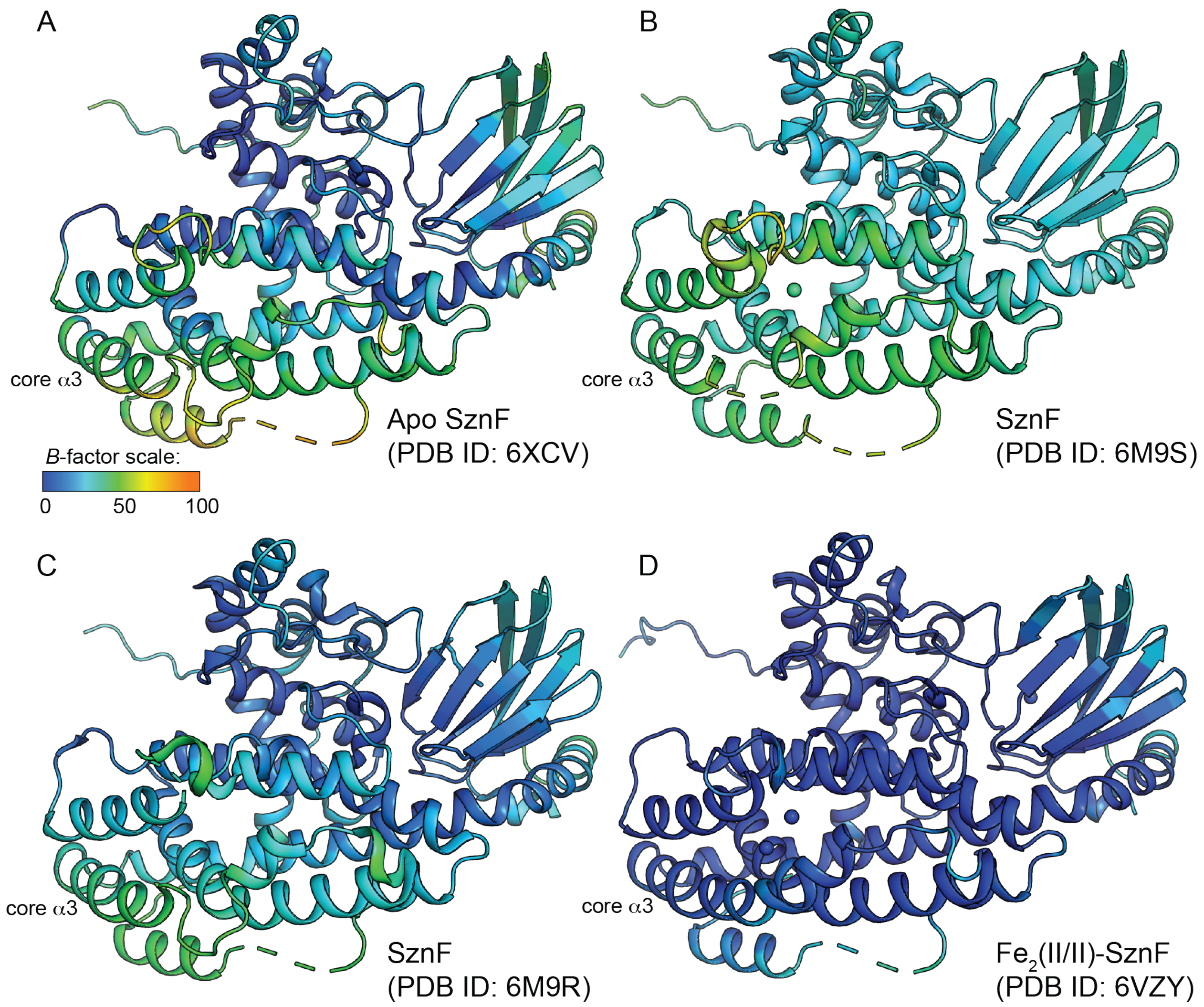


**Fig. S12.** Analysis of *B*-factors in SznF structures solved in previous work and this study. While core α3 remains flexible (highest *B*-factors) relative to other parts of the structure in all four models, the overall values of the *B*-factors are diminished in the HDO domain after Fe(II) is bound (right panel).

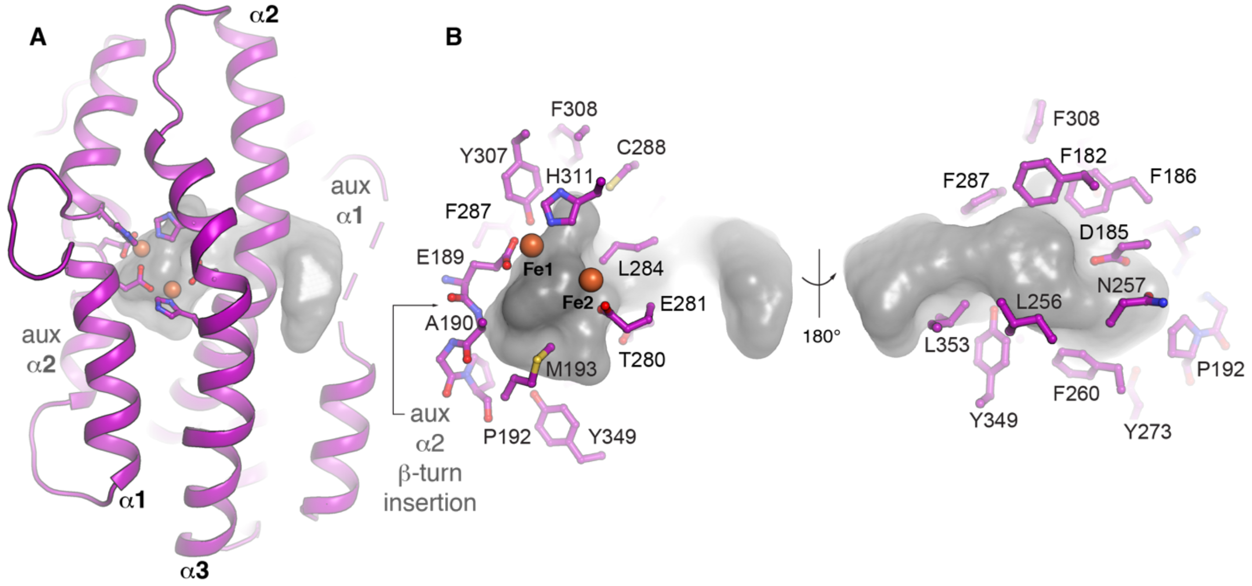


**Fig. S13.** Cavity mapping analysis of the HDO domain in $Fe_2$(II/II)-SznF. (*A*) A >1000 Å$^3$ cavity, mapped by the program HOLLOW (12) and analyzed by the DogSite Scorer server (49), extends from a disordered region (dashed line) in aux α1 near the surface of the protein to the HDO domain cofactor site, suggesting a plausible access route for the substrates. (*B*) Analysis of selected amino acid side chains lining the cavity shows that hydrophobic residues predominate.

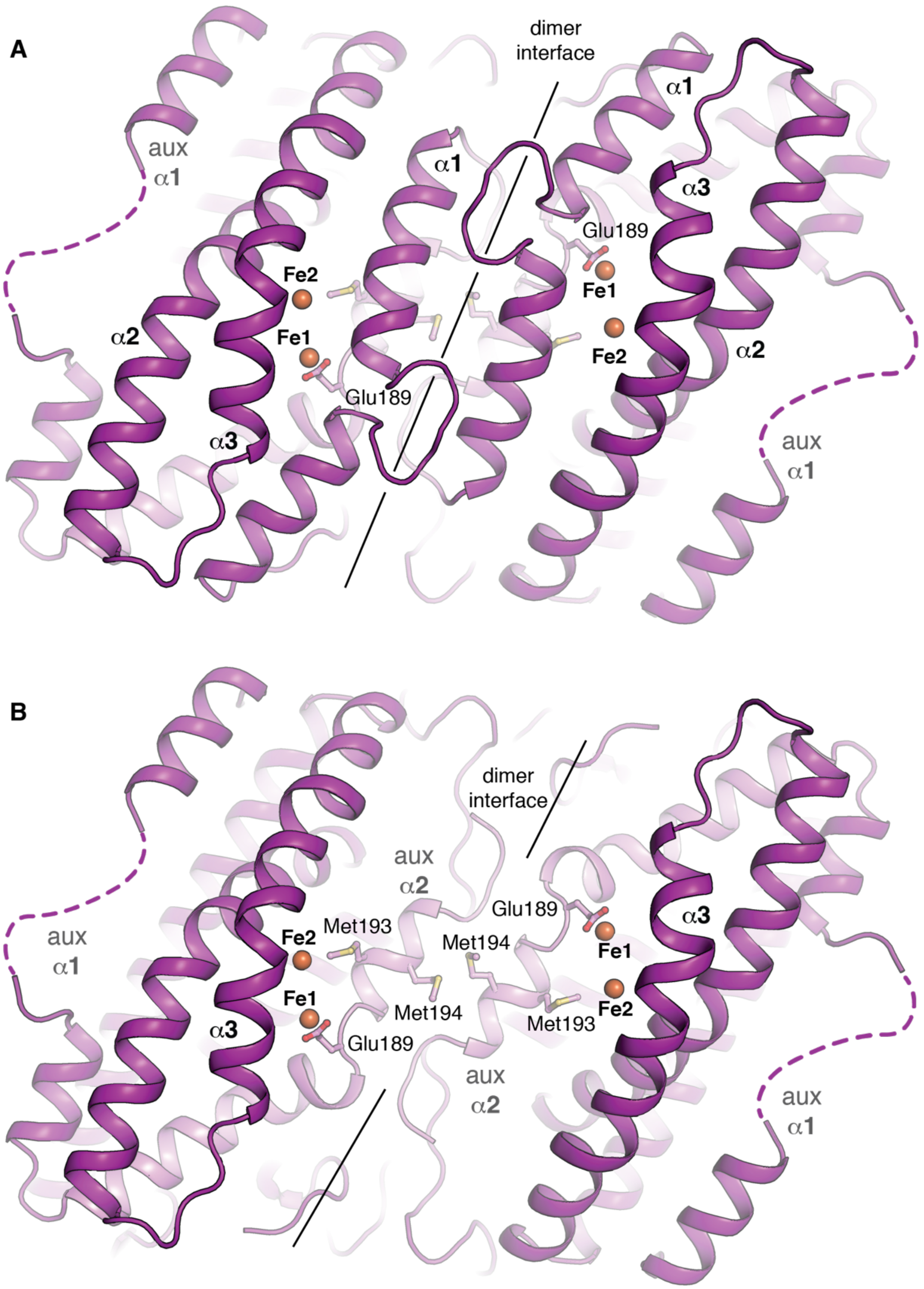


**Fig. S14.** Aux α2 and core α1 form symmetrical interactions across the SznF dimer interface that could communicate metal acquisition status in the other monomer. (*A*) A view of the dimer interface showing contacts in the core α1 interface. (*B*) A cutaway view showing the internal interface contacts between aux α2 structures. A pair of Met residues downstream of SznF ligand Glu189 form steric contacts at the dimer interface.

*Fe$_2$(II/II)-SznF•L-NMA models*

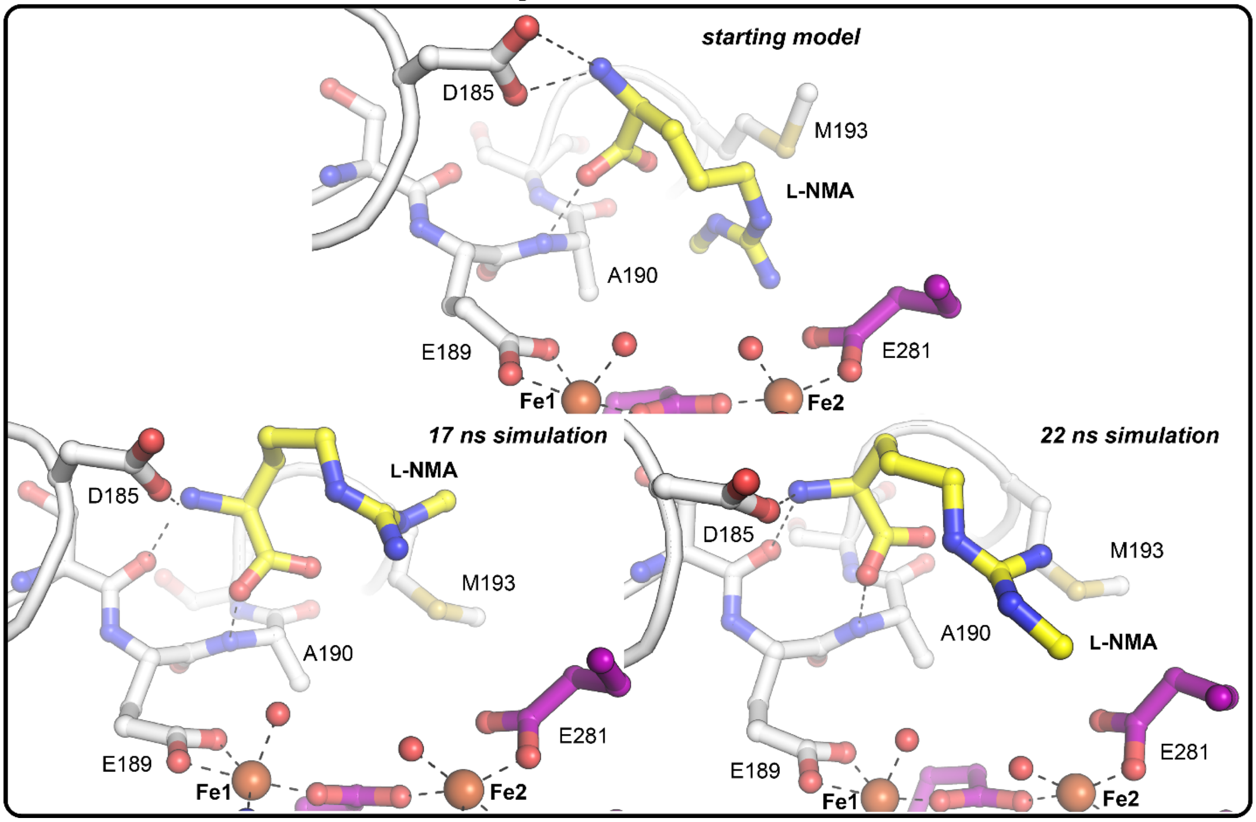


**Fig. S15.** Additional comparisons of an energy minimized docking model of the SznF HDO domain active site with the substrate, L-NMA, bound before (*top view*) and after (*bottom right and left views*) molecular dynamics simulation. The substrate binding pocket allows for 180° rotation of the side chain to position the ω' methyl group either proximal or distal to aux α2 (white). Selected amino acids and substrates are shown in *stick format*. Fe(II) ions and water molecules are shown as *orange and red spheres*, respectively.

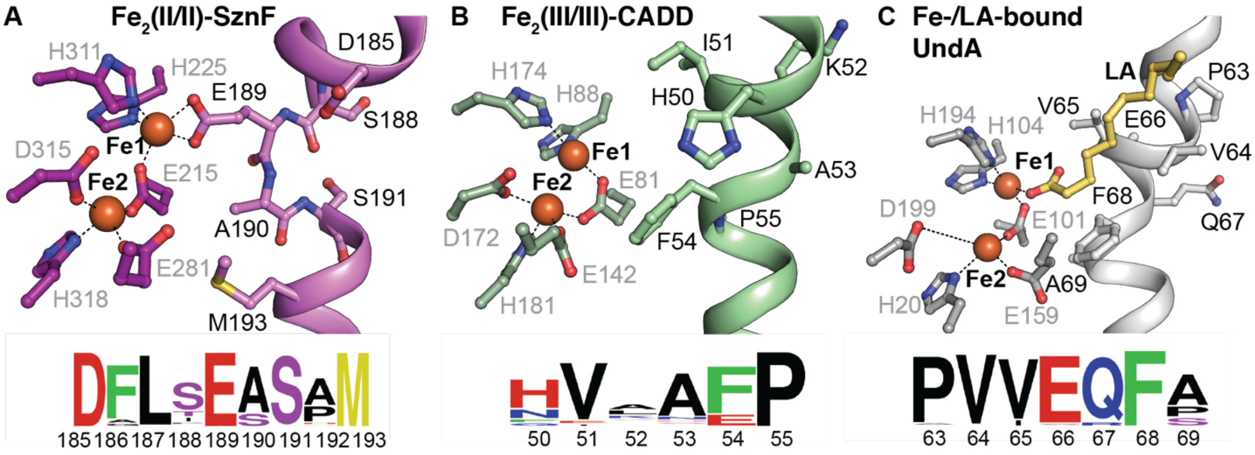


**Fig. S16.** Analysis of aux α2 structures in three HDOs, its importance in cofactor coordination in SznF by provision of the unexpected E189 ligand, and its possible role in substrate binding across the superfamily. (*A*) SznF (6VZY) aux α2 contains a β-turn motif that facilitates Fe1 coordination by providing Glu189 and provides additional polar contacts that are potentially important for substrate binding. The sequence motif (DFLSEAS) that enables this helical insertion and additional coordination interaction is highly conserved within the SznF sequence cluster (cluster 11 in the SSN shown in Fig. 5), as shown in the sequence logo diagram. The other characterized HDOs of divergent function including (*B*) CADD (PDB accession code 1RCW) and (*C*) UndA (PDB accession code 6P5Q) have uninterrupted aux α2 helices. CADD and UndA lack the additional Glu ligand and instead project hydrophobic residues into the active site. In UndA, these side chains form hydrophobic contacts with the alkyl chain of a coordinated lauric acid (LA) substrate. The carboxylate group of LA binds to Fe1 in the same location as Glu189 of SznF. Sequence logos for this region of aux α2 in the CADD and UndA SSN clusters (3 and 5, respectively, in the SSN shown in Fig. 5) show conservation of these sites.

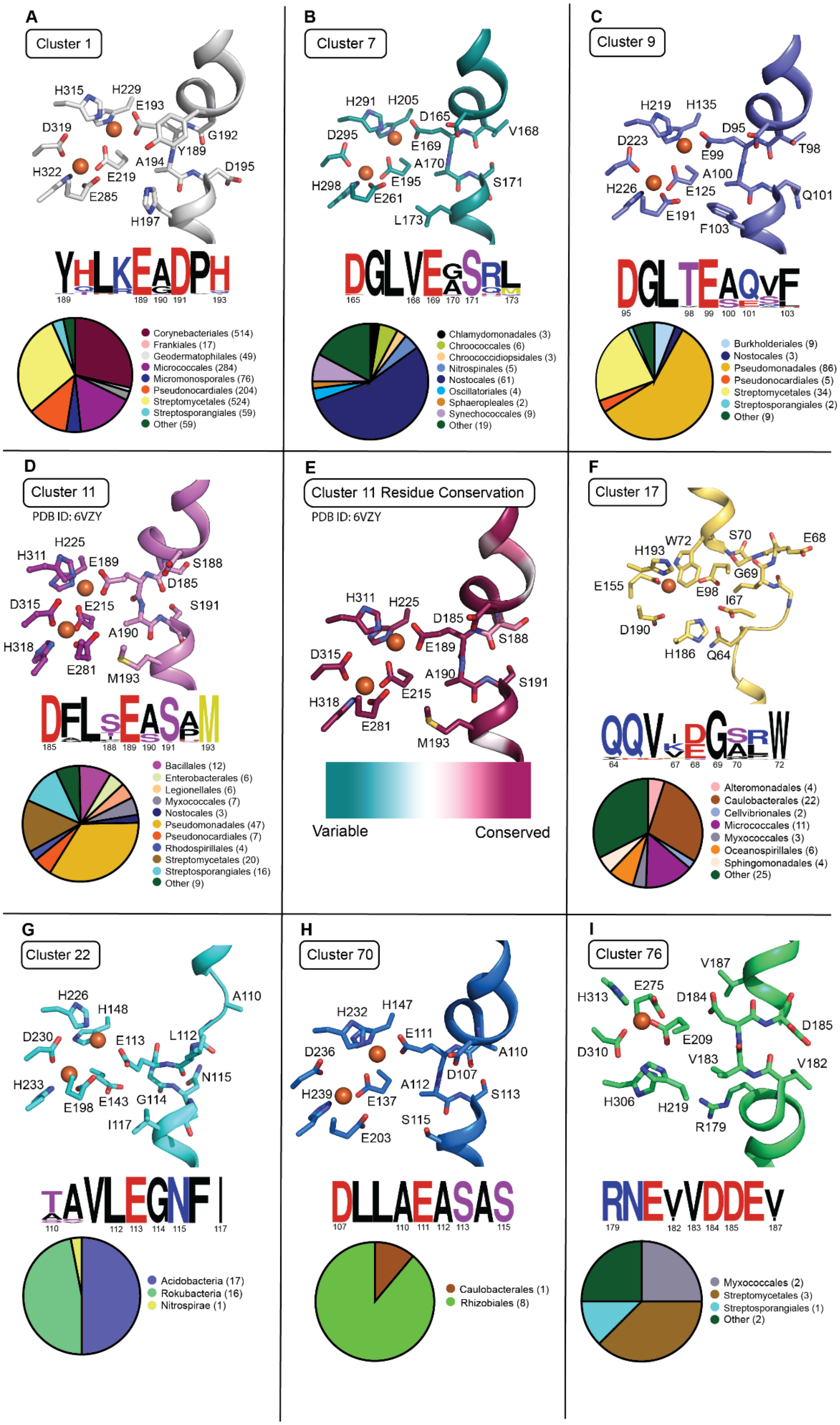

A
Cluster 1
H315
H229
E193
D319
G192
Y189
H322
A194
D195
E219
E285
H197
YHLKEADPH
189 189 190 191 193
Corynebacteriales (514)
Frankiales (17)
Geodermatophilales (49)
Micrococcales (284)
Micromonosporales (76)
Pseudonocardiales (204)
Streptomycetales (524)
Streptosporangiales (59)
Other (59)
B
Cluster 7
H291
H205
D165
D295
V168
E169
A170
E195
H298
S171
E261
L173
DGLVEGSRL
165 168 169 170 171 173
Chlamydomonadales (3)
Chroococcales (6)
Chroococcidiopsidales (3)
Nitrospinales (5)
Nostocales (61)
Oscillatoriales (4)
Sphaeropleales (2)
Synechococcales (9)
Other (19)
C
Cluster 9
H219
H135
D95
D223
T98
E99
A100
H226
E125
Q101
E191
F103
DGLTEAQVF
95 98 99 100 101 103
Burkholderiales (9)
Nostocales (3)
Pseudomonadales (86)
Pseudonocardiales (5)
Streptomycetales (34)
Streptosporangiales (2)
Other (9)
D
Cluster 11
PDB ID: 6VZY
H225
H311
E189
S188
D185
D315
E215
S191
A190
H318
E281
M193
DFLSEASAM
185 188 189 190 191 193
Bacillales (12)
Enterobacterales (6)
Legionellales (6)
Myxococcales (7)
Nostocales (3)
Pseudomonadales (47)
Pseudonocardiales (7)
Rhodospirillales (4)
Streptomycetales (20)
Streptosporangiales (16)
Other (9)
E
Cluster 11 Residue Conservation
PDB ID: 6VZY
H311
H225
D185
S188
E189
D315
A190
S191
E215
H318
E281
M193
Variable
Conserved
F
Cluster 17
H193
W72
S70
E68
G69
E155
E98
I67
D190
H186
Q64
QQVIDGSRW
64 67 68 69 70 72
Alteromonadales (4)
Caulobacterales (22)
Cellvibrionales (2)
Micrococcales (11)
Myxococcales (3)
Oceanospirillales (6)
Sphingomonadales (4)
Other (25)
G
Cluster 22
H226
H148
A110
D230
E113
L112
N115
H233
E198
E143
G114
I117
TAVLEGNFI
110 112 113 114 115 117
Acidobacteria (17)
Rokubacteria (16)
Nitrospirae (1)
H
Cluster 70
H232
H147
E111
D236
A110
D107
A112
H239
E137
S113
S115
E203
DLLAEASAS
107 110 111 112 113 115
Caulobacterales (1)
Rhizobiales (8)
I
Cluster 76
V187
E275
H313
D184
D185
E209
D310
V183
V182
H306
H219
R179
RNEVVDDEV
179 182 183 184 185 187
Myxococcales (2)
Streptomycetales (3)
Streptosporangiales (1)
Other (2)

**Fig. S17.** Homology models, sequence logos (50, 51), and biological distribution of sequences in SznF-like clusters from the SSN shown in Fig. 5. Homology models (*A*-*C*, *F*-*I*) were generated using SWISS-MODEL (52-57) with SznF (PDB accession code 6VZY) as a template. (*E*) SznF was colored according to sequence conservation using the ConSurf Database (13-15, 17, 58). Models indicate conservation of the β-turn motif within aux $\alpha 2$ across multiple clusters in the sequence similarity network (*A*-*E*, *H, I*). Models of sequences from clusters 17 and 22 (*F*, *G*) do not retain the β turn but conserve residues with sizes and properties similar to the corresponding residues in SznF. Sequence logo diagrams reveal conservation of the seventh carboxylate ligand directly upstream of a small residue within the β-turn. Pie charts highlight the representation of bacterial orders within (*A*) cluster 1 (n = 1,786), (*B*) cluster 7 (n = 112), (*C*) cluster 9 (n = 148), (*D*) cluster 11 (n = 137), (*F*) cluster 17 (n = 77), (*H*) cluster 70 (n = 9), and (*I*) cluster 76 (n = 8) with phyla listed in (*G*) cluster 22 (n = 34).

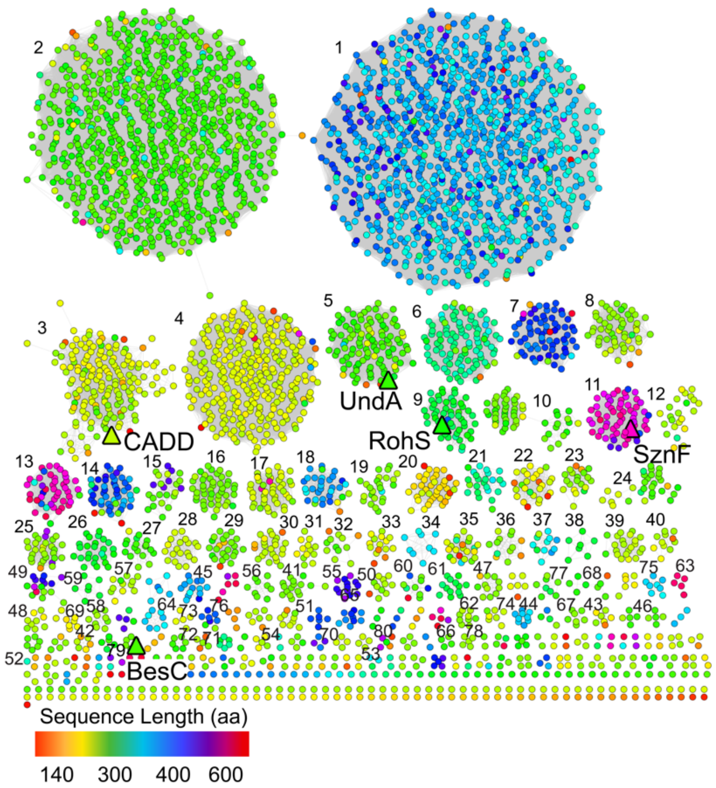


**Fig. S18.** SSN colored by sequence length. At an alignment score of 51, clusters generally contain sequences of similar lengths, indicating that sequences that cluster together may contain the same number of domains. Sequences in clusters 11 and 13 are of sufficient length to contain more than one domain.

**Table S1.** Data collection and refinement statistics for x-ray structures of SznF.

| | $Fe_2$(II/II)-SznF *native* | $Fe_2$(II/II)-SznF *Fe anomalous* | $Fe_2$(II/II)-SznF *Mn anomalous* | apo-SznF *native* |
|---|---|---|---|---|
| **Data collection** | | | | |
| Space group | $P\ 2_1\ 2_1\ 2_1$ | $P\ 2_1\ 2_1\ 2_1$ | $P\ 2_1\ 2_1\ 2_1$ | $P\ 2_1\ 2_1\ 2_1$ |
| Wavelength (Å) | 1.03319 | 1.73642 | *1.87848* | 1.03319 |
| Cell dimensions | | | | |
| *a*, *b*, *c* (Å) | 57.9, 105.8, 50.6 | 57.7, 105.5, 150.1 | 57.9, 105.7, 150.3 | 57.9, 108.1, 150.2 |
| $\alpha, \beta, \gamma$ (°) | 90.0, 90.0, 90.0 | 90.0, 90.0, 90.0 | 90.0, 90.0, 90.0 | 90.0, 90.0, 90.0 |
| Resolution (Å) | 50.00-1.66 (1.69-1.66) | 50.00-2.54 (2.58-2.54) | 50.00-2.74 (2.79-2.74) | 50.00-1.77 (1.80-1.77) |
| $R_{merge}$ | 0.074 (1.067) | 0.063 (0.291) | 0.104 (0.303) | 0.063 (0.835) |
| $R_{pim}$ | 0.033 (0.466) | 0.023 (0.110) | 0.022 (0.066) | 0.028 (0.379) |
| $I / \sigma I$ | 23.6 (2.1) | 34.5 (7.3) | 35 (10.1) | 26.5 (2.1) |
| $CC_{1/2}$ | 0.996 (0.677) | 1.00 (0.967) | 1.00 (0.978) | 0.937 (0.698) |
| Completeness (%) | 96.8 (100) | 100 (99.7) | 97.5 (91.2) | 90.6 (97.3) |
| Redundancy | 5.8 (6.1) | 8.9 (7.9) | 24.1 (19.8) | 5.7 (5.4) |
| **Refinement** | | | | |
| Resolution (Å) | 50.00-1.66 | | | 50.00-1.77 |
| No. reflections | 105729 | | | 82685 |
| $R_{work}$ / $R_{free}$ | 0.19/0.22 | | | 0.21/0.25 |
| No. atoms | | | | |
| Protein | 7499 | | | 7444 |
| Ligand/ion | 6 | | | |
| Water | 411 | | | 592 |
| *B*-factors | | | | |
| Protein | 18.0 | | | 26.5 |
| Ligand/ion | 14.1 | | | |
| Water | 19.7 | | | 29.4 |
| R.m.s. deviations | | | | |
| Bond lengths (Å) | 0.010 | | | 0.008 |
| Bond angles (°) | 1.076 | | | 0.930 |
| Molprobity clashscore | 1.84 ($99^{th}$ percentile) | | | 4.67 (97th percentile) |
| Rotamer outliers (%) | 0 | | | 0 |
| Ramachandran favored (%) | 99 | | | 98 |

*Values in parentheses are for highest-resolution shell.

**Table S2.** Characteristics of clusters deemed similar to SznF on the basis of conservation of the aux $\alpha$2 Glu side chain and putative β turn-forming residues.

| Cluster | Sequences | Ave. sequence length | Nodes | Sequences/node | Example genera |
|---|---|---|---|---|---|
| **1** | 1,786 | 342 | 1,009 | 1.77 | *Streptomyces*, *Mycobacterium*, *Rhodococcus* |
| **7** | 114 | 369 | 71 | 1.61 | *Nostoc*, *Nitrospina*, *Leptolyngbya* |
| **9** | 148 | 287 | 53 | 2.79 | *Pseudomonas, Streptomyces*, *Burkholderia* |
| **11** | 137 | 459 | 62 | 2.21 | *Pseudomonas*, *Actinomadura*, *Streptomyces* |
| **17** | 77 | 241 | 45 | 1.71 | *Brevundimos*, *Halomonas*, *Rheinheimera* |
| **22** | 34 | 220 | 27 | 1.26 | *Edaphobacter, Acidipila* |
| **70** | 9 | 381 | 4 | 2.25 | *Rhizobium, Phenylobacterium* |
| **76** | 8 | 340 | 8 | 1 | *Streptomyces, Actinomadura, Sorangium* |

**Table S3.** Accession numbers for reference sequences used for alignments in bioinformatics identification of HDO enzymes.

| ***Protein*** | ***NCBI accession ID*** |
|---|---|
| *SznF* | QBA82042.1 |
| *BesC* | WP_014151496.1 |
| *RohS* | WP_063774763.1 |
| *UndA* | WP_011062591.1 |
| *CADD* | O84616.1 |
